# Deficient spermiogenesis in mice lacking *Rlim*

**DOI:** 10.1101/2020.08.31.275248

**Authors:** Feng Wang, Maria G. Gervasi, Ana Bošković, Fengyun Sun, Vera D. Rinaldi, Jun Yu, Mary C. Wallingford, Darya A. Tourzani, Jesse Mager, Lihua J. Zhu, Oliver J. Rando, Pablo E. Visconti, Lara Strittmatter, Ingolf Bach

**Affiliations:** Department of Molecular, Cell and Cancer Biology, University of Massachusetts Medical School, Worcester, MA 01605; Department of Veterinary & Animal Sciences, University of Massachusetts Amherst, Amherst, MA 01003; Department of Biochemistry and Molecular Pharmacology, University of Massachusetts Medical School, Worcester, MA 01605; Program in Molecular Medicine, University of Massachusetts Medical School, Worcester, MA 01605; Program in Bioinformatics and Integrative Biology, University of Massachusetts Medical School, Worcester, MA 01605; Electron Microscopy Core, University of Massachusetts Medical School, Worcester, Massachusetts 01605

**Keywords:** Rlim, male reproduction, spermiogenesis, cytoplasmic reduction, epigenetic regulation, mouse genetics

## Abstract

The X-linked gene *Rlim* plays major roles in female mouse development and reproduction, where it is crucial for the maintenance of imprinted X chromosome inactivation in extraembryonic tissues of embryos. However, while females carrying a systemic *Rlim* knockout (KO) die around implantation, male *Rlim* KO mice appear healthy and are fertile, raising questions as to the pressures driving *Rlim* gene selection during evolution. Here we report an important role for *Rlim* in testis where it is highly expressed in post-meiotic round spermatids as well as in Sertoli cells. Systemic deletion of the *Rlim* gene leads to lower numbers of mature sperm that contains excess cytoplasm, leading to decreased sperm motility and *in vitro* fertilization rates. Targeting the conditional *Rlim* cKO specifically to the spermatogenic cell lineage largely recapitulates this phenotype. These results reveal functions of *Rlim* in male reproduction specifically in round spermatids during spermiogenesis with likely evolutionary implications.

## Introduction

In testes of adult animals, the differentiation of primordial germ cells (PGCs) during spermatogenesis occurs within seminiferous tubules. Spermiogenesis represents a late stage during spermatogenesis, in which post-meiotic round spermatids differentiate into mature spermatozoa by condensation of the spermatid DNA and formation of the sperm head, midpiece and tail (O’Donnell et al. 2011; O’Donnell 2014). Spermatozoa are then released into the lumen of seminiferous tubules, in a process called spermiation, which involves the remodeling and reduction of cytoplasm (Franca et al. 2016). Even though these processes are crucial for male reproduction they are poorly understood, and the number of genes involved remain limited.

The ubiquitin proteasome system (UPS) plays important roles in male reproduction (Richburg et al. 2014) with a testis-specific version of the proteasome complex (Kniepert and Groettrup 2014). Indeed, the ubiquitination of proteins in cells of the testis is required for functional spermatogenesis including spermiogenesis, and multiple steps during the progression of PGCs to mature spermatozoa critically depend on the UPS (Richburg et al. 2014). The UPS pathway is critically dependent upon E3 ubiquitin ligases, which provide substrate specificity by selecting target proteins for ubiquitination (Pickart 2001; Metzger et al. 2014).

The X-linked gene *Rlim* (also known as *Rnf12*) encodes a RING H2 type E3 ligase (Bach et al. 1999; Joazeiro and Weissman 2000; Metzger et al. 2014). While *Rlim* mRNA is widely expressed in many organs and cell types, RLIM protein is more selectively detected (Bach et al. 1999; Ostendorff et al. 2006). In cells RLIM shuttles between the cytoplasm and nucleus. Nuclear translocation is regulated by phosphorylation, and in many cell types RLIM is primarily detected in the nucleus (Jiao et al. 2013), where it controls not only levels and dynamics of various proteins and protein complexes involved in transcriptional regulation (Bach et al. 1999; Ostendorff et al. 2002; Kramer et al. 2003; Gungor et al. 2007; Her and Chung 2009; Johnsen et al. 2009; Huang et al. 2011; Gontan et al. 2012; Wang et al. 2019), but also its own expression via autoubiquitination (Ostendorff et al. 2002). In female mice, *Rlim* functions as a major epigenetic regulator of nurturing tissues. RLIM promotes the survival of milk-producing alveolar cells in mammary glands of pregnant and lactating females (Jiao et al. 2012). Moreover, RLIM is crucial for imprinted X chromosome inactivation (iXCI) (Shin et al. 2010; Wang et al. 2016; Gontan et al. 2018), the epigenetic silencing of one X chromosome in placental trophoblast cells early during female embryogenesis to achieve X dosage compensation (Payer 2016). Indeed, due to inhibited placental trophoblast development, the deletion of a maternally inherited *Rlim* allele results in early peri-implantation lethality specifically of females (Shin et al. 2010; Wang et al. 2016). In contrast, males systemically lacking *Rlim* appear to develop normally, are born at Mendelian ratios and are fertile as adults (Shin et al. 2010).

Here, we report high and dynamic *Rlim* mRNA and protein expression in the testis of male mice, where expression of RLIM protein is highly detected in round spermatids during spermiogenesis as well as in Sertoli cells. Indeed, we show that *Rlim* is required for the generation of normal sperm numbers with normal sperm cytoplasmic volume. Even though Sertoli cells are known to regulate cytoplasmic reduction in spermatozoa (O’Donnell et al. 2011; O’Donnell 2014), our genetic analyses reveal that this activity is mediated by *Rlim* expressed in the spermatogenic cell lineage. These results assign important functions of *Rlim* during spermiogenesis.

## Results

### Rlim expression in testis is highly regulated

To investigate potential functions of *Rlim* in mice in addition to XCI, we examined mRNA expression in various tissues isolated from adult mice via Northern blots. *Rlim* mRNA was detected in many tissues with highest levels in testis (Fig. 1A), consistent with published results (Bach et al. 1999; Ostendorff et al. 2000). However, while the *Rlim*-encoding mRNA in most tissues migrated around 7.5kb, a variant band at 2.4kb was detected in testis. Based on published RNA-seq data sets on mouse testes isolated at various post-partum stages (Margolin et al. 2014), mapping of reads to the *Rlim* locus revealed relatively homogenously distribution over all exons in 6-20 days post-partum (dpp) animals. However, most of the reads in sexually mature (38 dpp) animals mapped in exonic regions upstream of the TGA Stop codon, encompassing the 5’ non-coding region and the entire open reading frame (ORF), while most of the 3’ noncoding region was underrepresented (Fig. 1B). Indeed, consistent with the length of the observed variant *Rlim* mRNA, closer examination the *Rlim* cDNA sequence revealed a consensus alternative polyadenylation site (Proudfoot 2011) starting 69 bp downstream of the TGA Stop codon (Fig. 1C). Thus, in mature mouse testes a short, variant *Rlim* RNA is generated by alternative polyadenylation (Tian and Manley 2017).

**Fig. 1.**
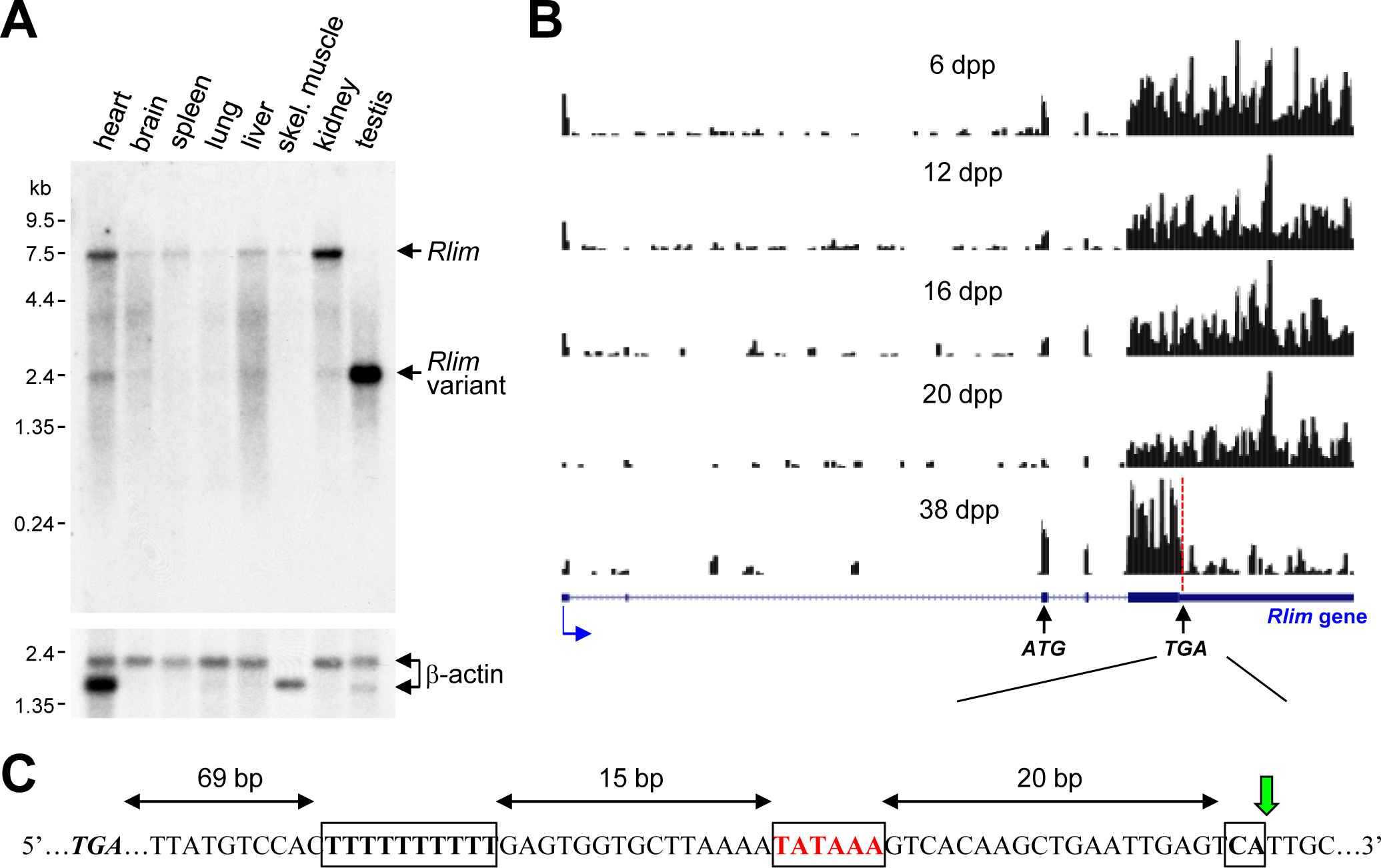
A short *Rlim* mRNA variant highly expressed in testis is generated via alternative polyadenylation during the maturation of male mice. **A)** A Northern blot containing RNA extracts from various adult mouse tissues (WT for *Rlim*) was hybridized with an *Rlim* probe (upper panel) and a probe recognizing β-actin as loading control (lower panel). **B)** Modified from the UCSC Genome Browser: Cumulative raw reads from RNA-seq datasets of testes RNA isolated from post-natal mice at 6, 12, 16, 20 and 38 days post-partum (dpp) (Margolin et al. 2014) were mapped on the *Rlim* locus (variable scales). Structure of the *Rlim* gene is shown below in blue with boxed exon regions. Protein coding regions are indicated in thicker stroke. Blue arrow indicates direction of transcription. ATG start codon, TGA stop codon and site of alternative polyadenylation sequence (red dotted line) is indicated. Note low relative read density 3’ of the alternative polyadenylation site specifically in 38 dpp animals. **C)** Nucleotide sequence containing an alternative polyadenylation site downstream of the TGA stop codon. Conserved motifs including a T-rich sequence, A/TATAAA and CA motifs are boxed. The cleavage position is indicated (green arrow).

Because expression of *Rlim* mRNA and protein can be strikingly different (Ostendorff et al. 2006), we examined RLIM protein in male reproductive tissues via immunohistochemistry (IHC) using an established RLIM antibody (Ostendorff et al. 2002; Ostendorff et al. 2006). Indeed, in testes we detected strong immunoreactivity in specific regions of some seminiferous tubules, representing differentiating spermatogenic cells (Fig. 2A). Moreover, we detected single RLIM-positive cells at the periphery of all tubules. As expected, little to no RLIM staining was detected in males carrying a Sox2-Cre (Hayashi et al. 2002) – mediated conditional knockout of the *Rlim* gene (cKO/Y^Sox2-Cre^), as these animals lack RLIM in somatic tissues as well as the germline (Shin et al. 2014; Wang et al. 2016). Because high RLIM levels appeared to be expressed at specific stages during spermatogenesis (Fig. 2A), we performed co-staining using an antibody against peanut agglutinin (PNA) that stains acrosomal structures allowing staging of spermatogenic cells within seminiferous tubules (Oakberg 1956b; Oakberg 1956a; Kotaja et al. 2004). Around seminiferous stage III, RLIM levels are detectable but low in all spermatogenic cells (Fig. 2B). However, indicated by RLIM/PNA co-staining, RLIM levels are dramatically upregulated specifically in round spermatids that have undergone meiosis at stages VI / VII, the timepoint when spermiogenesis begins (O’Donnell 2014; Qian et al. 2014). However, RLIM protein is no longer detectable in spermatozoa that are released during spermiation (O’Donnell et al. 2011) (Fig. 2B). We noted that the nuclei of RLIM-positive cells located at the periphery of seminiferous tubules displayed a triangular shape characteristic for Sertoli cells (Suppl. Fig. 1A). To identify the RLIM-positive cell types (Fig. 2A), we performed IHC, co-staining with antibodies against RLIM and the Sertoli cell marker GATA1 (Yomogida et al. 1994). Positive co-staining identified these cells as Sertoli cells (Fig. 2C) assigning RLIM as a novel marker for this cell type. Moreover, hybridizing epididymal sections with antibodies against *Rlim*, we also detected expression of RLIM in nuclei of epididymal epithelial cells (Suppl. Fig. 1B). This dynamic and regulated expression of *Rlim* suggests roles during male reproduction.

**Fig. 2.**
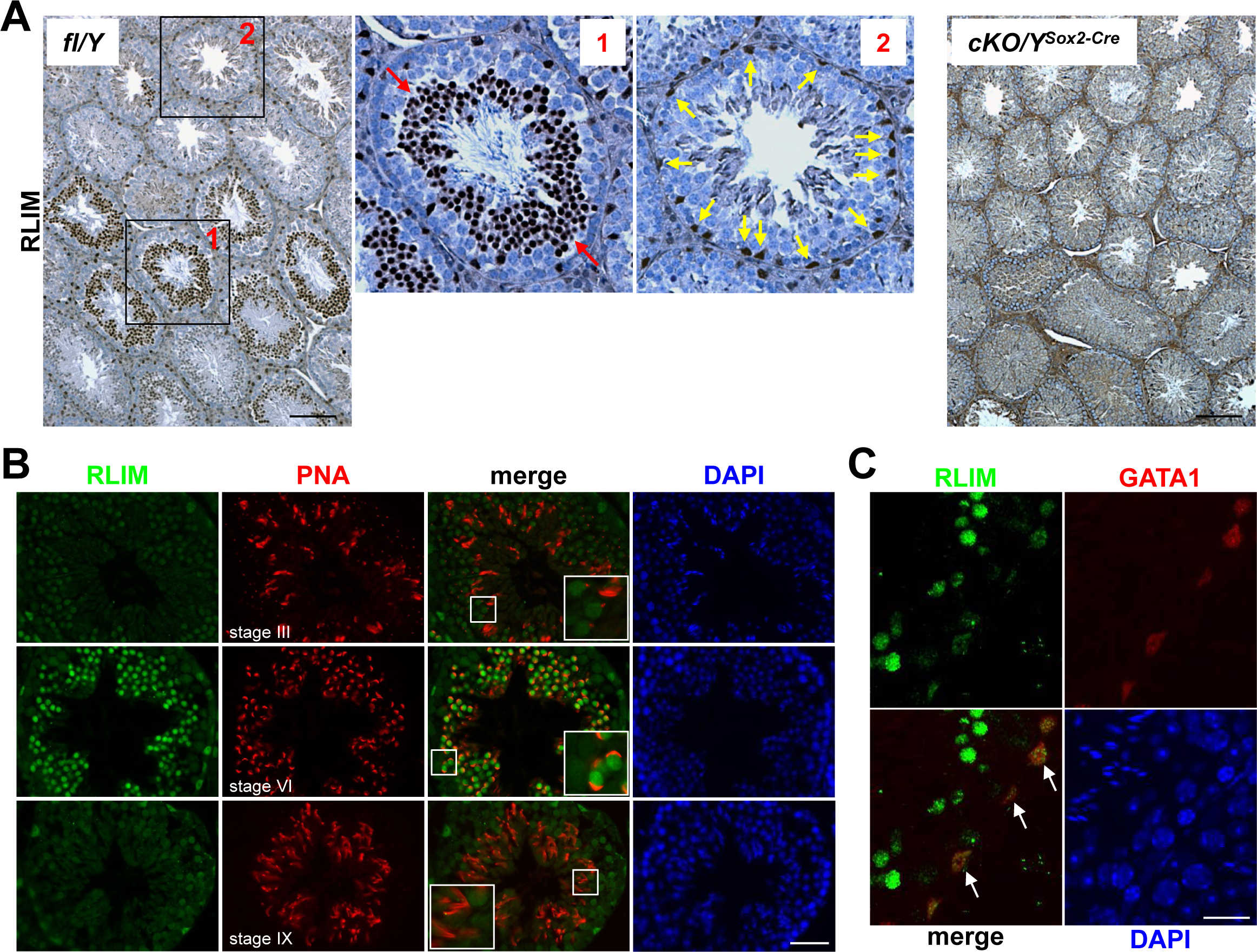
High RLIM protein expression specifically in round spermatids and in Sertoli cells. Tissue sections of mouse testes were stained using IHC with indicated antibodies. Boxed areas are shown in higher magnification. **A)** DAB staining of testes sections generated from fl/Y and, as negative control, *Rlim* cKO/Y^Sox2-Cre^ males littermates using antibodies against RLIM. Left panel: fl/Y male. Red arrows (box 1) and yellow arrows (box 2) point at spermatogenic cells and cells located in the periphery of seminiferous tubules that exhibit high RLIM staining, respectively. Right panel: cKO/Y male. Scale bar = 150 μm. **B)** IHC on fl/Y testis co-staining with antibodies against RLIM (green) and PNA (red) to determine differentiation stages of spermatogenic cells within seminiferous tubules (indicated). Scale bar = 60 μm. **C)** IHC on fl/Y testis co-staining with antibodies against RLIM (green) and GATA1 (red), a Sertoli cell marker. Scale bar = 25 μm.

### Diminished production and functionality of Rlim KO sperm

Next, we investigated functions of the *Rlim* gene in males using our conditional knockout (cKO) mouse model (Shin et al. 2010). Results on male mice systemically lacking *Rlim* induced by Cre recombinase transgenes (Shin et al. 2010; Shin et al. 2014; Wang et al. 2016) or by germline KO (Suppl. Fig. S2A) reveals that *Rlim* has no essential functions during male embryogenesis and post-natal development. To explore potential roles of the *Rlim* gene during male reproduction, we compared testes of 8 weeks-old animals systemically lacking *Rlim* either by a germline *Rlim* KO or with a Sox2-Cre – mediated *Rlim* cKO (cKO/Y^Sox2-Cre^) with fl/Y littermate controls. Indeed, in cKO/Y^Sox2-Cre^ and KO/Y mice the size and weight of the testis was significantly decreased when compared to fl/Y littermates, (Fig. 3A, B; Suppl. Fig. S2B). This was accompanied by lower numbers of mature sperm isolated in Caudal swim-out experiments (Fig. 3C; Suppl. Fig. S2C). In contrast, epididymal weights of cKO/Y^Sox2-Cre^ and fl/Y animals were similar (Suppl. Fig. S2D). Visualizing tubules that form the inner epididymal layers via de-lipidation (Tomer et al. 2014; Sylwestrak et al. 2016) revealed that the Cauda region of many cKO/Y^Sox2-Cre^ animals contained thinner tubules when compared to control males (Suppl. Fig. S2E)., consistent with decreased Caudal sperm (Fig. 3C). Moreover, cKO/Y^Sox2-Cre^ animals displayed a shortened Corpus region, while no obvious morphological phenotype was observed in the Caput region.

**Fig. 3.**
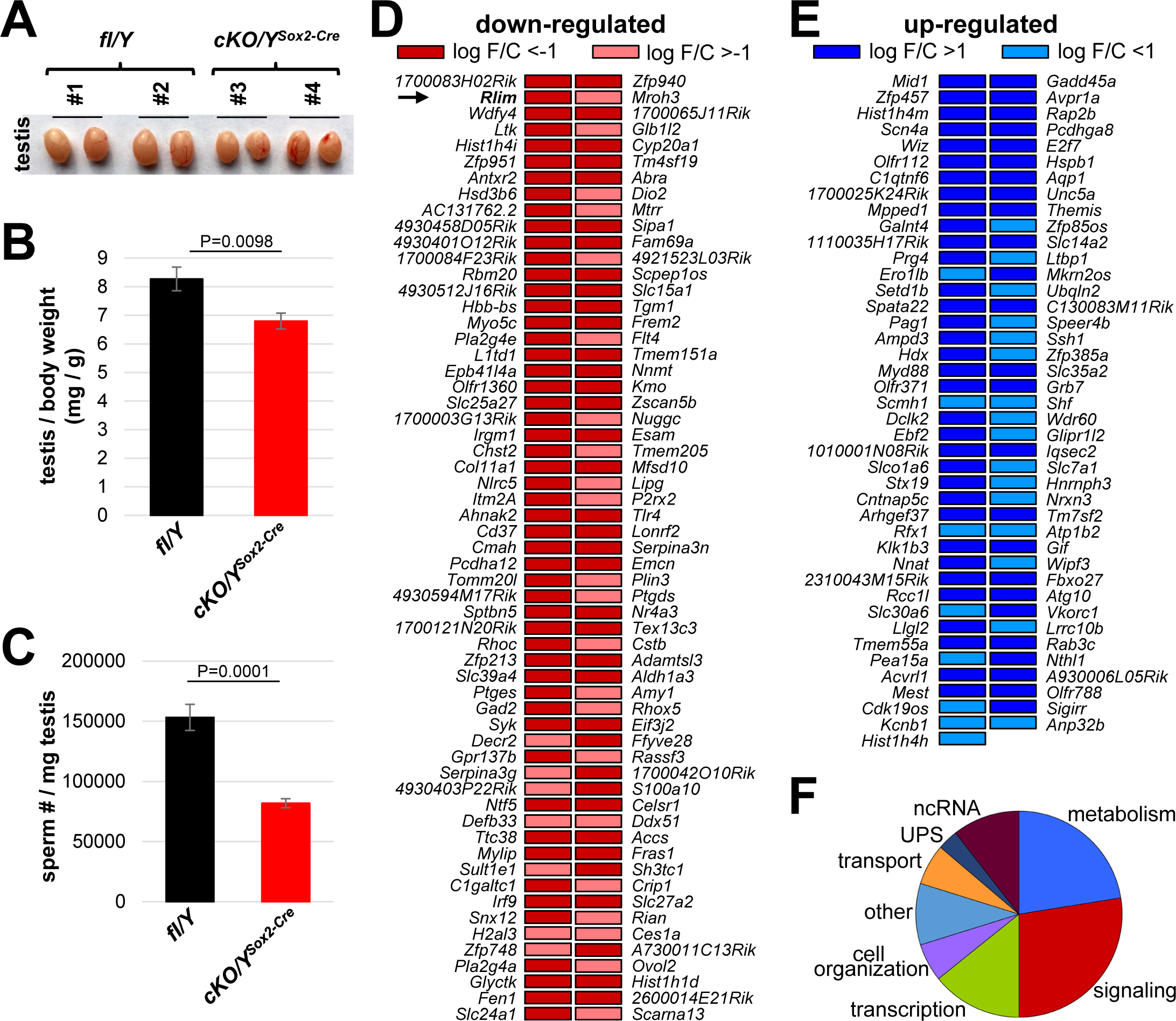
Lack of *Rlim* affects sperm production. Males systemically lacking *Rlim* were generated via Sox2-Cre mediated *Rlim* deletion (cKO/Y^Sox2-Cre^) and directly compared to fl/Y male littermates at 8 weeks of age. **A)** Deletion of *Rlim* results in smaller testes. Representative testes isolated from adult male fl/Y control animals (#1,2) and cKO/Y^Sox2-Cre^ littermates (#3,4) are shown. **B)** Significantly decreased weight of testes isolated from cKO/Y^Sox2-Cre^ animals (n=9) when compared to fl/Y littermates (n=7). Values were normalized against total body weight and represent the mean ± s.e.m. P values are shown (students t-test). **C)** Significantly decreased numbers of sperm in animals lacking *Rlim*. Cauda epididymal sperm were collected via swim-out in HTF medium. After 10 minutes of swim-out, total sperm numbers were determined (n=7 fl/Y; n=9 cKO/Y^Sox2-Cre^). s.e.m. and P values are indicated. **D, E)** Differentially expressed genes in testes of fl/Y and cKO/Y^Sox2-Cre^ mice as determined by RNA-seq experiments on biological replicates. Genes significantly (P<0.05) down-regulated and up-regulated upon the *Rlim* deletion in each experiment are shown in (C) and (D), respectively. Arrow indicates *Rlim*. **F)** Differentially expressed genes distribute in 8 functional categories that include metabolism, signaling, transcription, cell organization, transport, UPS, ncRNA and other.

As *Rlim* is a transcriptional regulator and the structural integrity of testes appeared to be intact in cKO/Y^Sox2-Cre^ males (Fig. 2A), we next examined effects of *Rlim* on genome wide gene expression. Thus, we performed RNA-seq experiments on RNA isolated from total testis of 8 weeks-old animals, comparing global gene expression in cKO/Y^Sox2-Cre^ and fl/Y male littermates. These experiments including library construction, sequencing and data processing were performed as previously described (Wang et al. 2017). Consistent with findings that attribute both positive and negative functions of RLIM for gene transcription (Bach et al. 1999; Gontan et al. 2012), statistical analyses revealed 118 down-regulated and 83 up-regulated genes (threshold P<0.05) in cKO/Y^Sox2-Cre^ animals (Figs. 3C, D). Functions of these genes fell mostly in 8 categories with around half of all differentially expressed genes involved in signaling (27.5%) and regulation of metabolism (22.5%), in particular regulatory functions on lipid metabolism (Fig. 3E). Other genes are involved in transcription/chromatin (14%), cell organization (6%), transport (6%) or the UPS (3%), while 9.5% of gene functions occupy multiple other cellular pathways. Around 10.5% of transcripts constituted non-coding (nc) RNAs. Combined, these results suggest functions of *Rlim* during spermatogenesis/sperm maturation.

Next, we investigated the characteristics and functionality of sperm isolated from the Cauda of 8 weeks-old males via swim-out. Indeed, cKO/Y^Sox2-Cre^ sperm displayed elevated rates of morphological abnormalities, including coiled midpieces and head malformations (Figs. 4A, B). However, the sperm capacitation-induced phosphorylation pathways, acrosomal status and induced acrosome reaction in *Rlim* KO sperm were similar to controls as judged by Western blot and PNA staining, respectively (Suppl. Fig. 3A-E), indicating that the deletion of *Rlim* did not affect general signaling. To examine sperm motility, we used CEROS computer-assisted semen analysis (CASA) in swim-out experiments at T0 and after 60 min (T60) under conditions that allow capacitation comparing cKO/Y^Sox2-Cre^ males with fl/Y littermates. The results revealed significant motility deficiencies of cKO/Y^Sox2-Cre^ sperm. Indeed, cKO/Y^Sox2-Cre^ sperm displayed decreased total motility, and out of the total motile sperm the percentages of sperm with progressive motility was lower, while the percentages of slow and weakly motile sperm populations were higher (Fig 4C-E). Thus, cKO/Y^Sox2-Cre^ males produce less sperm, which additionally is also less motile. Next, we tested for possible functional consequences of these deficiencies during *in vitro* fertilization (IVF) (Sharma et al. 2016), using oocytes originating from WT females and sperm isolated from either fl/Y or cKO/Y^Sox2-Cre^ littermates. In these experiments the numbers of oocytes and sperm cells were adjusted to 100-150 and 100,000, respectively, to achieve fertilization rates of around 80% for the control samples as judged by the number of embryos reaching cleavage stage 24h after adding sperm (Figs. 4F, G). The development of IVF embryos was monitored up to (96h) at which point blastocyst stage was reached by the majority of embryos generated by control sperm. Indeed, sperm isolated from cKO/Y^Sox2-Cre^ animals yielded in significantly less embryos reaching the appropriate developmental stage when compared to fl/Y littermate controls, both for cleavage and blastocyst stages (Fig. 4F, G). Combined, these data indicate that deletion of *Rlim* in male mice results in reduced sperm functionality.

**Fig. 4.**
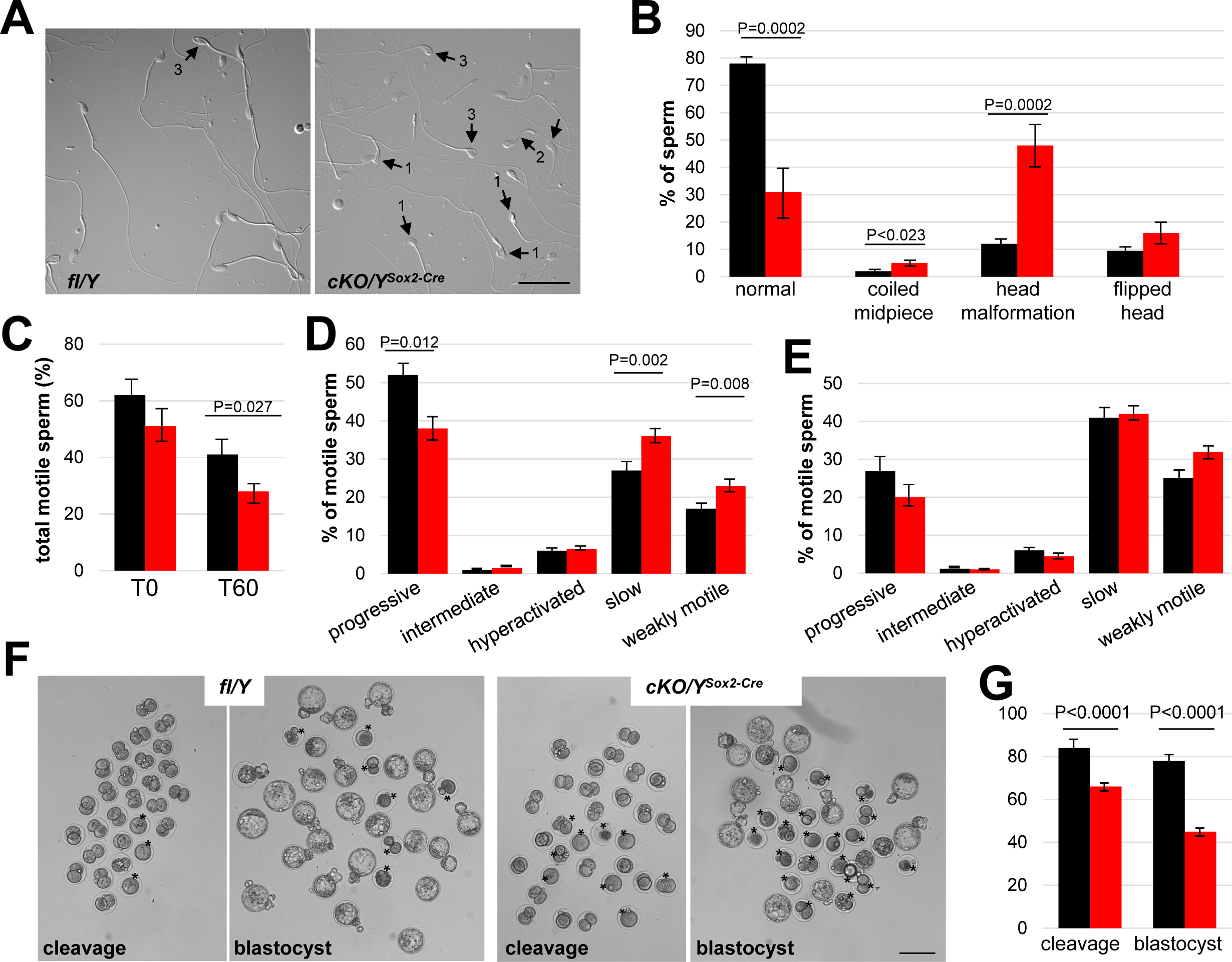
Increased abnormalities and decreased functionality in sperm lacking *Rlim*. Cauda epididymal sperm were collected via swim-out from 8 weeks-old males (n=7 fl/Y 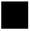; n=9 cKO/Y^Sox2-Cre^ 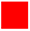) and morphology, motility and functionality were evaluated. **A)** Sperm morphology was assessed by light microscopy. Representative images of difference interference contrast (DIC) display the morphological patterns found indicated by arrows: 1, Head malformation; 2, Coiled midpiece; 3, Flipped head. Scale bar = 25 µm. **B)** Quantification of the morphology patterns. Percentages of normal sperm, coiled midpiece, head malformation, and flipped head out of the sperm population. At least 100 sperm per sample were counted, total sperm counted 1,146 for fl/Y and 1,077 for cKO/Y^Sox2-Cre^. Values represent the mean ± s.e.m. **C)** Sperm motility was evaluated in the swim-out (T0) and after 60 min of incubation in conditions that support capacitation (T60). Sperm motility was examined using the CEROS computer-assisted semen analysis (CASA) system. Percentage of total motile sperm in the population. n=12, values represent the mean ± s.e.m. **D)** Classification of type of motility (progressive, intermediate, hyperactivated, slow, and weakly motile; in percentage) out of the total motile population at T0. n=12, values represent the mean ± s.e.m. **E)** Classification of type of motility (in percentage) out of the total motile population after 60 minutes of incubation in capacitation conditions (T60). n=12, values represent the mean ± s.e.m. **F)** Representative images for cleavage and blastocyst stages at 24h and 96h, respectively. Asterisks indicate embryos not reaching anticipated embryonic stage. Scale bar = 100 μm. **G)** Summary of IVF results. n=142 and 313 presumed oocytes for fl/Y and cKO/Y sperm, respectively. Values represent the mean ± s.e.m.

### Increased size of cytoplasmic droplets in Rlim cKO sperm

Because of the decreased motility of *Rlim* cKO sperm (Figs. 4C-E), we examined the energetic status including amino acids, glycolysis, TCA cycle, pentose phosphate pathway, and nucleotide biosynthesis via metabolomic profiling of polar metabolites. We isolated 40-45 Mio cauda swim-out sperm each of 3 fl/Y and 5 cKO/Y^Sox2-Cre^ animals, and polar metabolites were measured using liquid chromatography coupled with mass spectrometry (LC-MS). Results showed that, unexpectedly, 66 metabolites were significantly more abundant in the cKO/Y^Sox2-Cre^ sperm, in particular levels of many amino acids, in contrast to only 5 metabolites that displayed lower levels (Fig. 5A). Analyses of these diverse metabolites did not yield in the identification of a specific energy production pathway but rather revealed that metabolite intermediates of many cellular pathways are increased in the *Rlim* cKO sperm, suggesting a more general problem. Thus, we included electron microscopy (EM) in our analyses. Interrogating sperm via scanning EM (SEM) revealed significantly more *Rlim* cKO/Y^Sox2-Cre^ sperm displaying cytoplasmic droplets in the midpiece, and these droplets were also increased in size as measured using ImageJ software (Figs. 5B-D). Because Cauda sperm has matured a considerable time in the epididymis that also expresses *Rlim* (Suppl. Fig. 1B), to distinguish testicular versus epididymal functions of *Rlim* in droplet formation we extended these studies to testicular sperm. Indeed, isolating testicular sperm from 30 fl/Y and cKO/Y^Sox2-Cre^ animals each, the numbers of isolated sperm per *Rlim* cKO testis were lower but this was no longer significant when compared to controls (Fig. 6A). We noted that the midpieces from cKO sperm were highly vulnerable to rupturing, while the prevalence of coiled midpieces appeared similar between cKO and control sperm (Fig. 6B). Moreover, we detected a low number of sperm that exhibited duplicated axonemes specifically in *Rlim* cKO/Y^Sox2-Cre^ but not in fl/Y sperm (Suppl. Fig. 4A). Again, the sizes of cytoplasmic droplets in testicular sperm were significantly increased (Fig. 6C, D). Transmission EM (TEM) on testes sections confirmed the occurrence of duplicated axonemes in cKO/Y^Sox2-Cre^ sperm as well as head malformations, while the overall structural integrity of the sperm tail and midpiece appeared normal (Suppl. Figs. 4A-D). Interestingly, the cytoplasmic pockets in the *Rlim* cKO sperm heads appeared more pronounced in sperm of the epididymal Caput region (Fig. 6E, F). These data reveal excessive cytoplasm in sperm heads and midpieces in males lacking *Rlim* and are consistent with testicular as opposed to epididymal functions of *Rlim*.

**Fig. 5.**
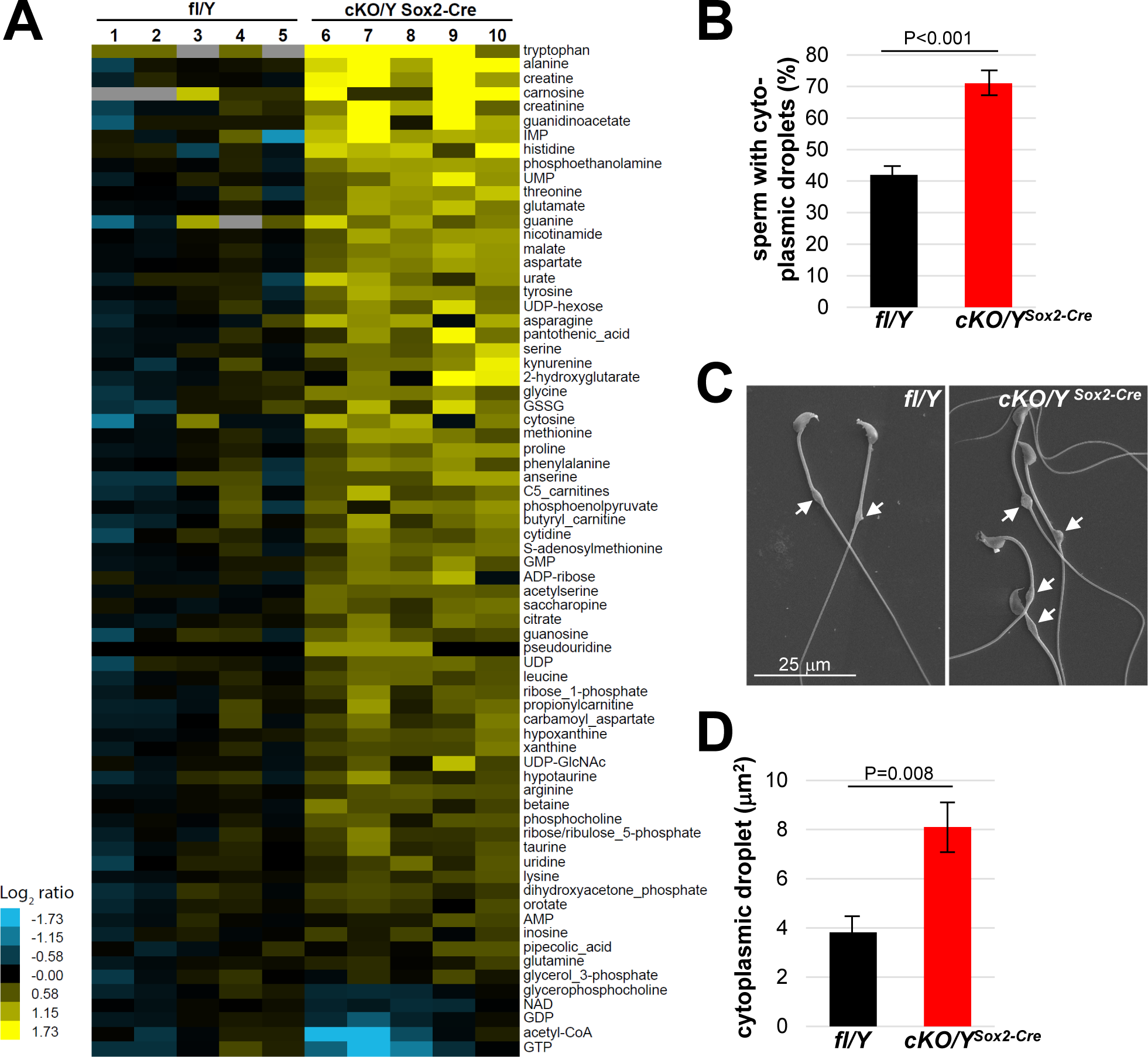
Increased size of cytoplasmic droplet in caudal sperm of males lacking *Rlim*. **A)** Cauda epididymal sperm were collected from 8 weeks-old fl/Y or cKO/Y^Sox2-Cre^ mice and polar metabolites were determined via LC-MS. Samples were run and data analyzed by the Metabolite Profiling Core Facility at the Whitehead Institute. Note general increased content of metabolites in cKO/Y^Sox2-Cre^ sperm. **B)** Cauda sperm was collected from 8 weeks-old fl/Y or cKO/Y^Sox2-Cre^ mice and after 10 min separation, immediately fixed in 2.5% glutaraldehyde followed by SEM analysis. Sperm with or without cytoplasmic droplets were counted. n=250, each. **C)** Increased size of cytoplasmic droplets in cKO/Y^Sox2-Cre^ sperm. Representative images are shown. Droplets are indicated by arrows. **D)** Increased size of cytoplasmic droplets in cKO/Y^Sox2-Cre^ sperm. Droplet surface size was determined via ImageJ. n=100, each.

**Fig. 6.**
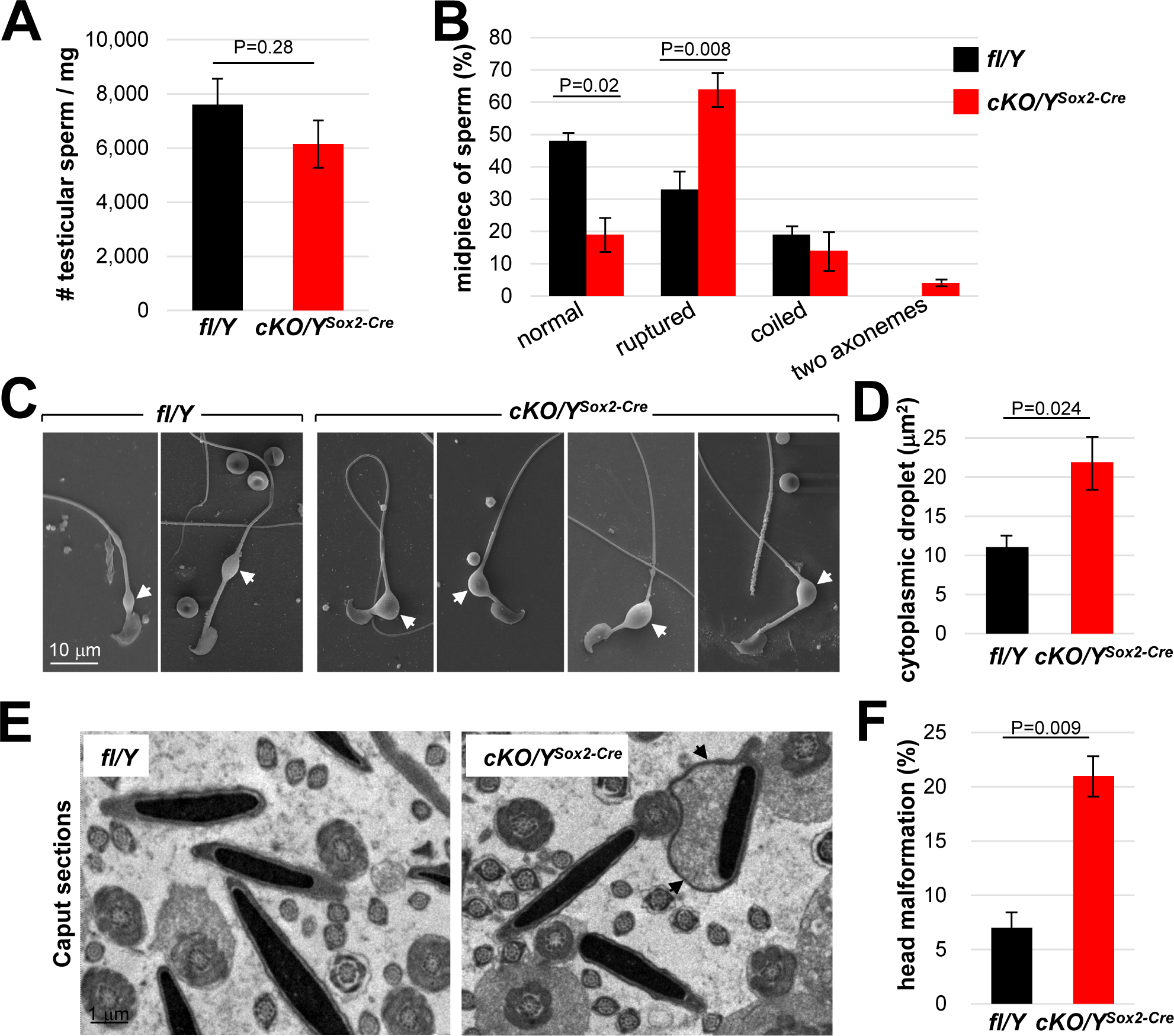
*Rlim* plays important roles for cytoplasmic reduction. Analyses of testicular sperm isolated from 8 weeks-old fl/Y or cKO/Y^Sox2-Cre^ mice. **A)** Quantification of sperm yield, normalized against testis weight. n=30, each genotype. **B)** Quantification of sperm morphology scoring ruptured, coiled sperm and sperm with two axonemes. n=250. **C)** Larger cytoplasmic droplets in cKO/Y^Sox2-Cre^ sperm. Representative images are shown. Droplets are indicated by arrows. **D)** Increased size of cytoplasmic droplets in cKO/Y^Sox2-Cre^ sperm. Droplet surface size was determined via ImageJ. n=100, each. **E)** Sperm head malformations within the epididymal Caput region as determined via TEM. Representative images are shown. Arrows point at cytoplasmic pocket. **F)** Quantification of sperm exhibiting excessive head cytoplasmic pocket.

### Functions of Rlim during spermiogenesis specifically in the spermatogenic cell lineage

Next, we addressed the question as to the cell type of *Rlim* action during spermiogenesis as RLIM protein is highly detectable both in Sertoli cells and in spermatogenic cell lineage specifically in round spermatids (Fig. 2). Because Sertoli cells play major roles in the regulation of spermiogenesis/spermiation (O’Donnell et al. 2011; O’Donnell 2014; Franca et al. 2016) and cell numbers are correlated with sperm production capacity (Griswold 1995), we elucidated number of cells positive for Sertoli cell marker GATA4, as in adult mice GATA4 expression does not vary with the cycle of the seminiferous epithelium (Yomogida et al. 1994). Counting GATA4-positive cells within seminiferous tubules, IHC revealed similar numbers of Sertoli cells in testes with or without *Rlim* (Suppl. Figs. 5A, B), indicating that *Rlim* is not required for Sertoli cell development and differentiation. Moreover, analyzing specific Sertoli cell structures involved in spermiation in testis sections via TEM, we did not detect major structural deficiencies in cells lacking *Rlim* including the formation of apical ectoplasmic specialization (ES) or the apical tubulobulbar complex (TBC) (Suppl. Figs. 5C, D).

To genetically elucidate the cell identity of *Rlim* function, we targeted the *Rlim* cKO via *Ngn3-Cre* (Schonhoff et al. 2004) to the spermatogenic cell lineage (Yoshida et al. 2004) and via *Sf1-Cre* (Dhillon et al. 2006) to Sertoli cells (Kim et al. 2007). Because of a mixed background of these Cre-driver mouse lines, cKO males were compared to their respective littermate controls. Indeed, *Rlim* cKO/Y^Ngn3-Cre^ mice displayed decreased testis weights and numbers of mature sperm isolated in caudal swim out experiments (Figs. 7A, B, respectively), recapitulating much of the phenotype observed in cKO/Y^Sox2-Cre^ animals that systemically lack *Rlim* (Fig. 3A, B). In contrast, no significant effects on testes weights and sperm numbers were measured in cKO/Y^Sf1-Cre^ animals. Consistent with these findings, SEM analyses of Caudal swim-out sperm revealed significantly increased numbers of sperm containing cytoplasmic droplets in cKO/Y^Ngn3-Cre^ animals when compared with cKO/Y^Sf1-Cre^ animals and their respective littermate controls (Fig. 7C). Moreover, focusing on sperm with cytoplasmic droplets, the droplet sizes in cKO/Y^Ngn3-Cre^ sperm were increased (Figs. 7D, E), similar to those of cKO/Y^Sox2-Cre^ sperm (Figs 5C, D). Combined, these results provide strong evidence that the upregulated expression of *Rlim* in round spermatids (Figs. 1, 2) plays important functions for the cytoplasmic remodeling during spermiogenesis/spermiation.

**Fig. 7.**
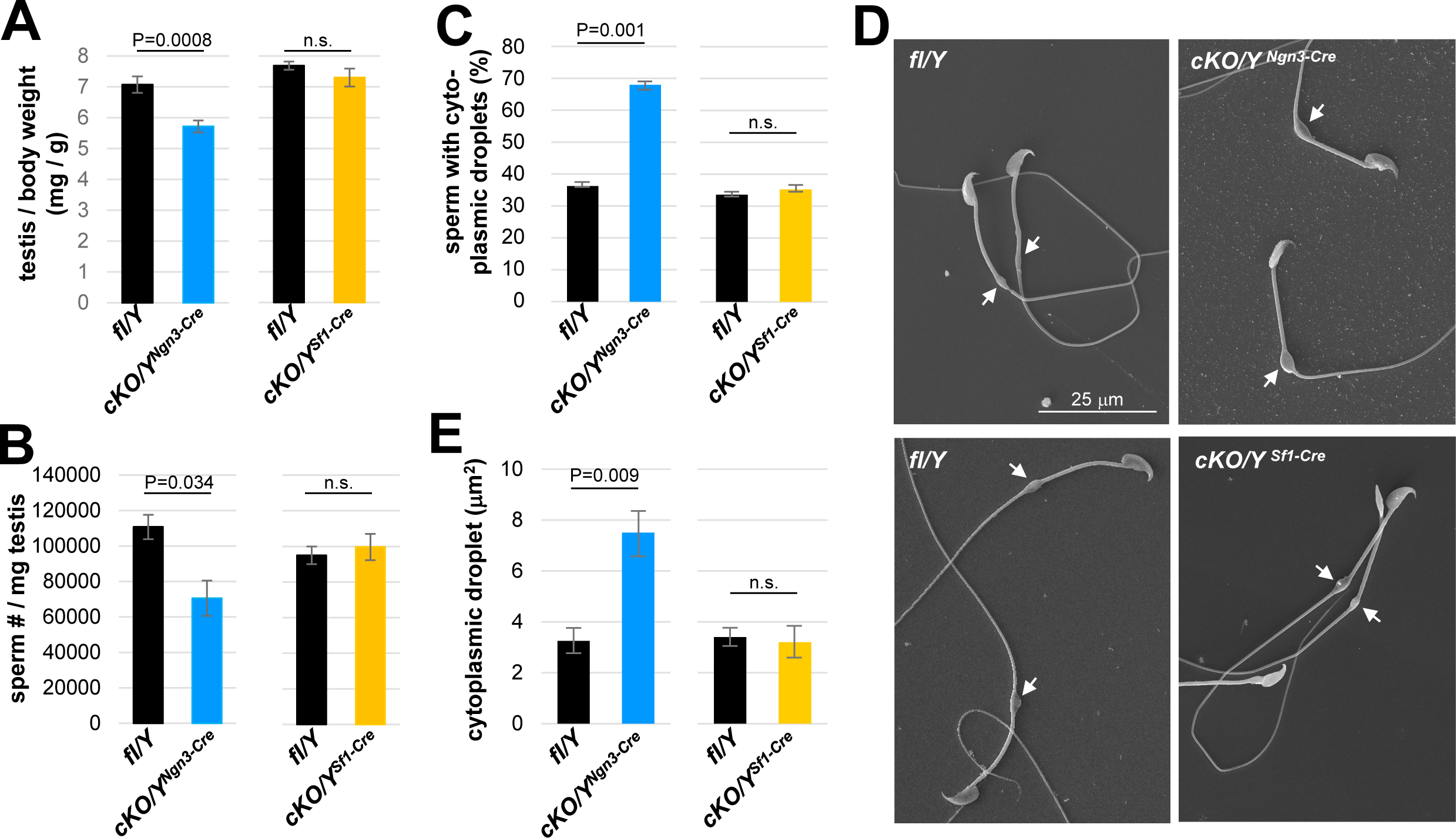
Functions of *Rlim* predominantly in the spermatogenic cell lineage. Animals with an *Rlim* cKO in the the spermatogenic cell lineage or in Sertoli cells were generated via Ngn3-Cre and SF1-Cre (cKO/Y^Ngn3-Cre^ and cKO/Y^SF1-Cre^), respectively, and directly compared to their fl/Y male littermates at 8 weeks of age. **A)** Significantly decreased weight of testes isolated from cKO/Y^Ngn3-Cre^ males but not from cKO/Y^Sf1-Cre^ animals (n=18 fl/Y; 14 cKO/Y^Ngn3-Cre^) (n=16 fl/Y; 10 cKO/Y^Sf1-Cre^). Values were normalized against total body weight and represent the mean ± s.e.m. P values are shown (students t-test). **B)** Significantly decreased numbers of sperm isolated from cKO/Y^Ngn3-Cre^ males but not from cKO/Y^Sf1-Cre^ animals. Cauda epididymal sperm were collected via swim-out in HTF medium. After 10 minutes of swim-out, total sperm numbers were determined (n=7 fl/Y; n=9 cKO/Y^Ngn3-Cre^; n=9 fl/Y; n=11 cKO/Y^Sf1-Cre^). s.e.m. and P values are indicated. **C)** Cauda sperm was collected from 8 weeks-old mice and visualized via SEM (n=3 per genotype). Sperm with or without cytoplasmic droplets were counted. n=250, per animal. **D)** Increased size of cytoplasmic droplets in cKO/Y^Ngn3-Cre^ sperm. Representative SEM images are shown. Upper panels: cKO/Y^Ngn3-Cre^ and fl/Y control. Lower panel: cKO/Y^Sf1-Cre^ and fl/Y control. Droplets are indicated by arrows. **E)** Increased size of cytoplasmic droplets in cKO/Y^Ngn3-Cre^ sperm. Droplet surface size of SEM images was determined via ImageJ. n=100, each.

## Discussion

Our results reveal robust expression of *Rlim* in male reproductive organs, particularly in testis, where the expression pattern was highly dynamic both at the mRNA and protein levels. While somatic tissues predominantly express a long 7.5 kb *Rlim* encoding mRNA, in testis, a short alternative 2.4 kb *Rlim* mRNA is expressed that barely covers the full ORF (Fig. 1). The appearance of this variant mRNA in males coincides with sexual maturation, suggesting that this form is pre-dominantly expressed in differentiating round spermatids, which express high levels of RLIM protein (Fig. 2A). Interestingly, mapping the reads of single embryo RNA-seq data to the *Rlim* locus reveals the variant mRNA as the predominant *Rlim* mRNA species also at preimplantation stages (Wang et al. 2016), when RLIM is highly expressed and exerts its crucial functions during iXCI in female mice (Shin et al. 2010). Thus, the expression of the short variant *Rlim* mRNA correlates with elevated RLIM protein levels and, because *Rlim* acts in the spermatogenic cell lineage (Fig. 7), also coincides with the exertion of its *in vivo* function in males. Indeed, alternative mRNAs display different time of synthesis, mRNA stability and/or translational efficiency (Tian and Manley 2017). At the protein level we find RLIM expression in spermatogenic cells is upregulated from low levels at stage III to high levels peaking in post-meiotic round spermatids at stage VI / VII. As this upregulation occurs at a stage during spermatogenic differentiation when meiotic sex chromosome inactivation (MSCI) has occurred (Turner 2007; Turner 2015), *Rlim* joins a number of genes that escape MSCI (Namekawa et al. 2006; Mueller et al. 2008; Berletch et al. 2010; Sin et al. 2012) and it is tempting to speculate that this escape is connected with the occurrence of the short, alternative *Rlim* mRNA. Moreover, even though *Rlim* is X-linked, the finding that the RLIM protein is detected in most/all round spermatids, including those that presumably harbor a Y chromosome, is explained by the fact that cytoplasmic bridges exist between spermatids and that RLIM efficiently shuttles between nuclei in heterokaryon cells (Jiao et al. 2013).

Sperm produced by males lacking *Rlim* is dysfunctional with decreased motility and increased cytoplasm and head abnormalities. Because excess cytoplasm affects sperm motility, morphology including head morphology as well as fertilization potential (Cooper 2011; Rengan et al. 2012), and the cytoplasmic volume is regulated during spermiogenesis just after RLIM protein is highly detected in spermatids, it is likely that the increased cytoplasmic volume is responsible for much of the defects detected in sperm lacking *Rlim*. Moreover, as the midpiece cytoplasm is particularly important for sperm osmoregulation (Cooper 2011; Rengan et al. 2012), an increased size of cytoplasmic droplets is predicted to render sperm more vulnerable to osmotic challenges, which may ultimately lead to midpiece rupturing (Fig. 6). In this context, an increased sperm cytoplasm has been associated with higher activities of specific enzymes of the energy pathway including G6PDH (Aitken et al. 1994; Yuan et al. 2013), which diverts glucose metabolism away from glycolysis towards the pentose pathway. It is thus tempting to speculate that increased G6PDH activity might be partially responsible for the decreased Acetyl-CoA levels measured in sperm lacking *Rlim* (Fig. 5A). As we detected neither increased cell death (not shown) nor signs of changes in chromatin packaging in sperm heads, our combined data suggests that flawed cytoplasmic removal during spermiogenesis likely represents a major underlying deficiency in *Rlim* KO sperm.

Concerning the cell type of *Rlim* action, RLIM protein is detected in spermatogenic cells specifically in round spermatids, in Sertoli cells and in epididymal epithelial cells. Our combined data reveals that lack of RLIM specifically in round spermatids is responsible for much of the observed sperm phenotype. This is demonstrated by targeting the *Rlim* cKO via Ngn3-Cre in the spermatogenic cell lineage, resulting in defective spermiogenesis, and similar sperm phenotypes when compared to the systemic *Rlim* cKO (Figs 3, 7). Moreover, the dramatic upregulation of *Rlim* in round spermatids coincides with the beginning of spermiogenesis. In contrast, targeting the *Rlim* cKO via Sf1-Cre to Sertoli cells failed to induce a testis/sperm phenotype and together with our data that Sertoli cell number and appearance seem normal, these results provide strong evidence that *Rlim* in Sertoli cells is not involved in the regulation of spermiogenesis provided by this cell type. Concerning functions of *Rlim* in the epididymis, as mentioned above, targeting the cKO via Ngn-cre yields in similar sperm phenotypes as a systemic KO. Moreover, testicular sperm from *Rlim* cKO/Y^Sox2-Cre^ animals exhibit head malformations and increased cytoplasmic droplet sizes (Fig. 6), indicating that lack of *Rlim* in epididymis is not involved in these deficiencies. Because Caudal sperm yields are significantly decreased in animals lacking *Rlim*, it is likely that the changes in Cauda and Corpus size observed in *Rlim* cKO/Y^Sox2-Cre^ animals (Suppl. Fig. 2E) are a consequence of less sperm content in this organ. Thus, while we cannot exclude minor functions of *Rlim* in Sertoli cells and epididymal epithelial cells, our results provide strong evidence that *Rlim* in round spermatids is required for normal spermiogenesis. Therefore, *Rlim* adds to a very limited number of genes in the spermatogenic cell lineage that regulate the cytoplasmic reduction during spermiogenesis (Zheng et al. 2007; Mikolcevic et al. 2012).

Considering early lethality of female mouse embryos with a maternally transmitted *Rlim* mutation (Shin et al. 2010), mathematical modeling of this exclusively female phenotype indicates that a deleterious mutation in the *Rlim* gene would shift selective evolutionary pressure entirely on females leading to a gender bias towards males in a mouse population over time (Jiao et al. 2012). Because gender biases in mouse populations represent an unfavorable strategy for reproduction (Hamilton 1967), it is likely that the observed functions of *Rlim* during male reproduction will contribute to counter-act gender biases induced by its female function in mouse populations in the wild. However, the extend of evolutionary gender balancing is unclear at this stage, as this function depends on mating behavior and the integration of multiple evolutionary drivers.

Combined, our data provide first evidence that in addition to crucial epigenetic functions in female embryogenesis and reproduction, the E3 ubiquitin ligase RLIM also occupies important roles during the reproduction of male mice. These results have major implications for epigenetic regulation and the evolution of *Rlim* and emphasize the importance of the UPS in male reproduction.

## Material and Methods

### Mice

Mice used in this study and genotyping have been described; *Rlim fl/fl* (Shin et al. 2010), *Sox2-Cre* (JAX #008454; in C57BL/6) (Hayashi et al. 2002; Shin et al. 2014), *Ngn3-Cre* (kind gift of A. Leiter; JAX #005667; mixed background); (Schonhoff et al. 2004) and *Sf1-Cre* (JAX #012462; mixed background) (Dhillon et al. 2006). *Rlim* mice were bred and maintained in a C57BL/6 background. All mice were housed in the animal facility of UMMS and utilized according to NIH guidelines and those established by the UMMS Institute of Animal Care and Usage Committee (IACUC).

### Antibodies, IHC and Northern blotting

Primary antibodies used for immunostainings were rabbit RLIM (Ostendorff et al. 2002), Lectin PNA from Arachis hypogaea (peanut), GATA1 (Santa Cruz Biotechnology, sc265), GATA4 (Abcam, ab84593), pPKAs (Cell Signaling, # 9624), PY (Millipore, clone G10), Alexa Fluor 488-conjugated lectin peanut agglutinin (ThermoFisher, # L21409). Secondary antibodies were Alexa Fluor® 488 Donkey Anti-Rabbit IgG (Invitrogen, A21206), Alexa Fluor® 488 Goat Anti-mouse IgG (Invitrogen, A11029), Alexa Fluor® 568 Goat Anti-Rabbit IgG (Invitrogen, A11011), Alexa Fluor® 568 Goat Anti-rat IgG (Invitrogen, A11077), and Alexa Fluor® 568 Goat Anti-mouse IgG (Invitrogen, A11004). Immunohistochemical (co-)staining on paraffin-embedded tissue sections were carried out as previously reported (Shin et al. 2010). Northern blot analysis was performed on a membrane containing RNA from adult mouse tissues (Clontech). As probe, we used a 568 bp SacI fragment isolated from mouse *Rlim* cDNA, which was ^32^P-labeled via random priming (Gibco BRL) as previously described (Ostendorff et al. 2000).

### RNA-seq and data analyses

RNA-seq on RNA isolated from testes of fl/Y and cKO/Y males including library construction and sequencing on a NextSeq 500 was carried out essentially as described (Wang et al. 2016; Wang et al. 2017). Reads (paired end 35 bp) were aligned to the mouse genome (mm10) using TopHat (version 2.0.12) (Trapnell et al. 2009), with default setting except set parameter read-mismatches was set to 2, followed by running HTSeq (version 0.6.1p1) (Anders et al. 2015), Bioconductor packages edgeR (version 3.10.0) (Robinson and Smyth 2007; Robinson et al. 2010) and ChIPpeakAnno (version 3.2.0) (Zhu et al. 2010; Zhu 2013) for transcriptome quantification, differential gene expression analysis, and annotation. For edgeR, we followed the workflow as described in (Anders et al. 2013).

### De-lipidation of epididymis

De-lipidation of epididymis was performed based on previously described protocols (Tomer et al. 2014; Sylwestrak et al. 2016). Briefly, isolated organs were fixed with 4% paraformaldehyde (PFA) in PBS for 32 hours at room temperature (RT), then rinsed with PBS for three times of at least two hours. Tissues were kept at 4°C in PBS with 0.02% sodium azide until the time of tissue processing. In order to visualize tubules that form the inner layers of the epididymis, a de-lipidation step was performed. De-lipidation was done passively by incubating the organ with 4% SDS/PBS at RT in an orbital shaker for two weeks. The 4% SDS/PBS solution was changed every other day. At the end of the second week the organ was rinsed with PBS for three times of least four hours and finally placed on a refractive index matching solution (RIMS: 0.17M iodixanol; 0.4M diatrizoic acid, 1M n-methyl-d-glucamine, 0.01% sodium azide), 24 hours prior to imaging.

### Collection of sperm

Epididymal Cauda sperm collected via swim-out and testicular sperm which were analyzed in this study was collected from 8 weeks-old fl/Y or cKO/Y mice. Briefly, Cauda epididymides were dissected and placed in 1 ml of modified Krebs-Ringer medium (m-TYH; 100 mM NaCl, 4.7 mM KCl, 1.2 mM KH_2_PO_4_, 1.2 mM MgSO_4_, 5.5 mM Glucose, 0.8 mM Pyruvic Acid, 1.7 mM Calcium Chloride, 20 mM HEPES). Sperm was allowed to swim-out for 10 min at 37 °C and then the epididymides were removed. Concentration of all sperm was calculated using a Neubauer hemocytometer. For mature testicular spermatozoa isolation, testes from one 8 weeks old mouse were minced in a 35-mm Petri dish containing 1 ml 150 mM NaCl. Finely minced tissue slurry was then transferred to a 15ml conical tube and set aside for 3-5 minutes to allow tissue pieces to settle down. Next, the cell suspension was loaded onto 10.5 ml of 52% isotonic percoll (Sigma). The tubes were then centrifuged at 15000 x g for 10 minutes at 10°C. The pellet was resuspended in 10 ml of 150 mM NaCl and spun at 900 x g at 4°C for 10 minutes followed by three washes with 150 mM NaCl at 4000 x g at 4°C for 5 min.

### Sperm analyses

Concerning the assessment of sperm morphology, after swim-out, sperm suspensions (50 µl) were fixed with paraformaldehyde 4 % (w/v; EMS, Hatfield, MA) in phosphate buffered saline (PBS) for 10 min at room temperature. Then samples were centrifuged at 800 x g for 5 min, washed with PBS, and let air-dry on poly-L-lysine-coated glass slides. Samples were mounted using VectaShield mounting media (H-1000, Vector Laboratories, Burlingame, CA) and differential interference contrast (DIC) images were taken using a 60X objective (Nikon, PlanApo, NA 1.49) in an inverted microscope (Nikon Eclipse TE300). Images were analyzed using the freeware ImageJ v1.52e (http://imagej.nih.gov/ij/download.html), and sperm morphology was classified into the following categories: normal, coiled midpiece, head malformation, flipped head, and residual cytoplasmic droplet. Results are shown as percentages, at least 200 sperm per sample were counted in single-blinded experiments. Concerning the analysis of sperm acrosomal status, sperm from the swim-out suspension were loaded into glass slides and let air-dry for 15 min. After that samples were fixed, and the acrosomes were stained as described below in the sperm acrosome reaction section. Fluorescence and phase contrast images were taken in a Nikon Eclipse TE300 fluorescence microscope using a 40X objective (Nikon). Images were analyzed using the freeware ImageJ v1.52e, and at least 200 sperm per sample were counted in single-blinded experiments. Results are shown as percentage of acrosome intact sperm.

Concerning sperm capacitation and analysis of swim-out cauda sperm motility, sperm were incubated at 37 °C for 60 min in m-TYH (Non-Cap) or in m-TYH supplemented with 15 mM NaHCO_3_ and 5 mg/ml BSA (Cap). Sperm motility was evaluated in the swim out (T=0) and after 60 min of incubation in capacitation conditions (T=60). Briefly, sperm suspensions (30 µl) were loaded into pre-warmed chamber slides (Leja slides, Spectrum Technologies, Healdsburg, CA) and placed on a warmed microscope stage at 37 °C. Sperm motility was examined using the CEROS computer-assisted semen analysis (CASA) system (Hamilton Thorne Research, Beverly, MA). Acquisition parameters were set as follows: frames acquired: 90; frame rate: 60 Hz; minimum cell size: 4 pixels; static head size: 0.13–2.43; static head intensity: 0.10–1.52; and static head elongation: 5–100. At least 5 microscopy fields corresponding to a minimum of 200 sperm were analyzed in each experiment. Data were analyzed using the CASAnova software (Goodson et al. 2011).

Concerning SDS-PAGE and Western blotting, swim-out cauda sperm samples were centrifuged at 12,000 x g for 2 min, washed in 1 ml of PBS, and then centrifuged at 12,100 ⨯ g for 3 minutes. Sperm proteins were extracted by resuspending the remaining pellets in Laemmli sample buffer (Laemmli 1970), boiled for 5 min and centrifuged once more at 12,100 ⨯ g for 5 min. Protein extracts (supernatant) were then supplemented with β-mercaptoethanol 5 % (v/v), and boiled again for 4 min. Protein extracts equivalent to 2.5 ⨯ 10^5^ sperm/lane were subjected to SDS–PAGE, and electro-transferred to PVDF membranes (Bio-Rad, Waltham, MA). PVDF membranes were blocked with 5 % (w/v) fat-free milk in tris buffered saline containing 0.1 % (v/v) Tween 20 (T-TBS) and immunoblotted with anti-pPKAs antibody (clone 100G7E, 1:10,000) overnight at 4°C to detect phosphorylated PKA substrates. Then, membranes were incubated with HRP-conjugated anti-rabbit secondary antibody diluted in T-TBS (1:10,000) for 60 min at room temperature. Detection was done with an enhanced chemiluminescence ECL plus kit (GE Healthcare) as per manufacturer instructions. After developing of pPKAs, membranes were stripped at 65 °C for 20 min in 2 % (w/v) SDS, 0.74 % (v/v) β-mercaptoethanol, 62.5 mM Tris (pH 6.5), blocked with fish gelatin 20 % (v/v; Sigma cat # G7765, St. Louis, MO) in T-TBS for 60 min at room temperature and re-blotted with anti-PY antibody (clone 4G10, 1:10000) to detect proteins phosphorylated in tyrosine residues. Membranes were then incubated with HRP-conjugated anti-mouse secondary antibody diluted in T-TBS (1:10,000) for 60 min at room temperature. Detection was done with an enhanced chemiluminescence ECL plus kit (GE Healthcare) as per manufacturer instructions.

Concerning swim-out cauda sperm acrosome reaction, after 60 min of capacitation in m-TYH Cap medium, sperm samples were incubated with progesterone (10 µM) or with DMSO (vehicle) in m-TYH Cap at 37 °C for 30 min. Then, sperm were loaded into glass slides and let air-dry for 15 min. Sperm were fixed by incubation with paraformaldehyde 4 % (w/v) in PBS at room temperature for 15 min, washed three times (5 min each) with PBS and permeabilized with 0.1 % (v/v) Triton X-100 for 3 min. After permeabilization, samples were washed three times with PBS, and then incubated with Alexa Fluor 488-conjugated lectin peanut agglutinin (PNA) in PBS at room temperature for 30 min. Before mounting with Vectashield (Vector Laboratories, Burlingame, CA), samples were washed three times with PBS for 5 min each time. Epifluorescence images were taken in a Nikon Eclipse TE300 fluorescence microscope using a 40X objective (Nikon). Phase contrast images were taken in parallel. Images were analyzed using the freeware ImageJ v1.52e, and at least 200 sperm per sample were counted in single-blinded experiments. Results are shown as percentage of acrosome reacted sperm.

### In Vitro Fertilization (IVF)

IVF experiments were performed essentially as described (Sharma et al. 2016) with some modifications. Briefly, female C57BL/6J mice (age 6–8 weeks) were superovulated by intraperitoneal injection of pregnant mare’s serum gonadotropin (PMSG; 5 U; Calbiochem) followed by human chorionic gonadotropin (hCG; 5 U; Sigma-Aldrich) 48 h later (Yamashita et al. 2008). Metaphase II-arrested oocytes tightly packed with cumulus cells were collected from the oviductal ampulla 14 h after hCG injection and placed in a 100-μl drop of human tubal fluid (HTF; Millipore) medium covered with mineral oil. Fresh cauda epididymal sperm of fl/Y and cKO/Y littermates (age, 2–4 mo) were swum-up in 1 ml HTF medium for 10 min and capacitated by incubation for another 30 min at 37°C under 5% CO_2_. An aliquot (1.0 × 10^5^ cells) of the capacitated sperm suspension was added to 100-μl drop of HTF medium containing the oocytes. After incubation at 37°C under 5% CO_2_ for 4 h, the presumed zygotes were washed with KSOM medium to remove cumulus cells, sperm and debris, and then incubated in a 50-μl drop of KSOM medium. *In vitro* fertilized embryos were analyzed at cleavage (24h) and blastocyst stages (96h).

### Metabolomic profiling

Polar metabolite measurements were performed by the Metabolite Profiling Core Facility at the Whitehead Institute for Biomedical Research (Cambridge, MA). For LC-MS analyses, 45 Mio epididymal cauda swim-out sperm were collected each from 5 cKO/Y^Sox2-Cre^ and 3 fl/y animals in extraction mix containing 80% methanol and isotopically labeled amino acids. Peak area ratios of each metabolite were determined and log_2_ ratio relative to the mean of control (fl/Y) sperm determined.

### Electron microscopy

SEM was performed on caudal epididymis sperm and sperm isolated from testes. Caudal sperm was let to dissociate in HTF medium and then fixed in 2.5% glutaraldehyde, 2% paraformaldehyde in 0.1M Na Cacodylate buffer. The fixed sperm was then placed on poly Lysine covered cover slips and left to adhere for 10 minutes. The samples were rinsed three times in the same fixation buffer, post fixed in aqueous 1% (w/v) OsO4 for 1hr at RT, dehydrated through a graded ethanol series to ethanol 100% (x3), and then they were critically point dried. The cover slips were then mounted on double sided carbon tape onto aluminum SEM stubs and grounded with colloidal silver paint, sputter coated with 12 nm of gold-palladium and were imaged using secondary electron (SEI) mode with a FEI Quanta 200 MKII FEG SEM.

TEM was performed on ultrathin sections on Caput epididymides and testes. Tissues were dissected and immediately immersed in 2.5% glutaraldehyde in 0.1 M Na Cacodylate buffer, pH 7.2. for 60 min at RT. The samples were rinsed three times in the same fixation buffer and post-fixed with 1% osmium tetroxide for 1h at room temperature. Samples were then washed three times with ddH_2_O for 10 minutes, and in block stained with a 1% Uranyl Acetate aqueous solution (w/v) at 6 °C overnight. After 3 rinses in ddH2O the samples were dehydrated through a graded ethanol series of 20% increments, before two changes in 100% ethanol. Samples were then infiltrated first with two changes of 100% Propylene Oxide and then with a 50%/50% propylene oxide / SPI-Pon 812 resin mixture. The following day five changes of fresh 100% SPI-Pon 812 resin were done before the samples were polymerized at 68°C in flat embedding molds. The samples were then reoriented, and thin sections (approx. 70 nm) were placed on copper support grids and contrasted with Lead citrate and Uranyl acetate. Sections were examined using the a CM10 TEM with 100Kv accelerating voltage, and images were captured using a Gatan TEM CCD camera.

### Data availability

RNA-seq data have been deposited with the Gene Expression Omnibus (GEO) database under accession number GSE114593.

## Acknowledgments

We are grateful to M. Krykbaeva, A. Schlueter, G. Pazour, J. Shin and E. Torres for advice and/or experimental help. I.B. is a member of the University of Massachusetts DERC (DK32520). This work was supported from NIH grants R01GM128168 to I.B., R01HD080224 and DP1ES025458 to O.J.R, and R01HD38082 to P.E.V.

## Autor contributions

Conceptualization: I.B. and F.W.; Methodology: F.W., M.G.G., A.B., V.D.R., J.Y.; Investigation, F.W., M.G.G., A.B., V.D.R., M.C.W., F.S., D.A.T., L.S., and I.B.; Supervision: I.B., L.S., P.E.V., O.J.R., J.M., and L.J.Z.; Writing: I.B., M.G.G., F.W.

## References

Aitken J, Krausz C, Buckingham D. 1994. Relationships between biochemical markers for residual sperm cytoplasm, reactive oxygen species generation, and the presence of leukocytes and precursor germ cells in human sperm suspensions. Molecular reproduction and development 39: 268–279.

Anders S, McCarthy DJ, Chen Y, Okoniewski M, Smyth GK, Huber W, Robinson MD. 2013. Count-based differential expression analysis of RNA sequencing data using R and Bioconductor. Nature protocols 8: 1765–1786.

Anders S, Pyl PT, Huber W. 2015. HTSeq--a Python framework to work with high-throughput sequencing data. Bioinformatics 31: 166–169.

Bach I, Rodriguez-Esteban C, Carriere C, Bhushan A, Krones A, Rose DW, Glass CK, Andersen B, Izpisua Belmonte JC, Rosenfeld MG. 1999. RLIM inhibits functional activity of LIM homeodomain transcription factors via recruitment of the histone deacetylase complex. Nature genetics 22: 394–399.

Berletch JB, Yang F, Disteche CM. 2010. Escape from X inactivation in mice and humans. Genome biology 11: 213.

Cooper TG. 2011. The epididymis, cytoplasmic droplets and male fertility. Asian J Androl 13: 130–138.

Dhillon H, Zigman JM, Ye C, Lee CE, McGovern RA, Tang V, Kenny CD, Christiansen LM, White RD, Edelstein EA et al. 2006. Leptin directly activates SF1 neurons in the VMH, and this action by leptin is required for normal body-weight homeostasis. Neuron 49: 191–203.

Franca LR, Hess RA, Dufour JM, Hofmann MC, Griswold MD. 2016. The Sertoli cell: one hundred fifty years of beauty and plasticity. Andrology 4: 189–212.

Gontan C, Achame EM, Demmers J, Barakat TS, Rentmeester E, van IW, Grootegoed JA, Gribnau J. 2012. RNF12 initiates X-chromosome inactivation by targeting REX1 for degradation. Nature 485: 386–390.

Gontan C, Mira-Bontenbal H, Magaraki A, Dupont C, Barakat TS, Rentmeester E, Demmers J, Gribnau J. 2018. REX1 is the critical target of RNF12 in imprinted X chromosome inactivation in mice. Nat Commun 9: 4752.

Goodson SG, Zhang Z, Tsuruta JK, Wang W, O’Brien DA. 2011. Classification of mouse sperm motility patterns using an automated multiclass support vector machines model. Biology of reproduction 84: 1207–1215.

Griswold MD. 1995. Interactions between germ cells and Sertoli cells in the testis. Biology of reproduction 52: 211–216.

Gungor C, Taniguchi-Ishigaki N, Ma H, Drung A, Tursun B, Ostendorff HP, Bossenz M, Becker CG, Becker T, Bach I. 2007. Proteasomal selection of multiprotein complexes recruited by LIM homeodomain transcription factors. Proceedings of the National Academy of Sciences of the United States of America 104: 15000–15005.

Hamilton WD. 1967. Extraordinary sex ratios. A sex-ratio theory for sex linkage and inbreeding has new implications in cytogenetics and entomology. Science 156: 477–488.

Hayashi S, Lewis P, Pevny L, McMahon AP. 2002. Efficient gene modulation in mouse epiblast using a Sox2Cre transgenic mouse strain. Mechanisms of development 119 Suppl 1: S97–S101.

Her YR, Chung IK. 2009. Ubiquitin Ligase RLIM Modulates Telomere Length Homeostasis through a Proteolysis of TRF1. The Journal of biological chemistry 284: 8557–8566.

Huang Y, Yang Y, Gao R, Yang X, Yan X, Wang C, Jiang S, Yu L. 2011. RLIM interacts with Smurf2 and promotes TGF-beta induced U2OS cell migration. Biochemical and biophysical research communications 414: 181–185.

Jiao B, Ma H, Shokhirev MN, Drung A, Yang Q, Shin J, Lu S, Byron M, Kalantry S, Mercurio AM et al. 2012. Paternal RLIM/Rnf12 is a survival factor for milk-producing alveolar cells. Cell 149: 630–641.

Jiao B, Taniguchi-Ishigaki N, Gungor C, Peters MA, Chen YW, Riethdorf S, Drung A, Ahronian LG, Shin J, Pagnis R et al. 2013. Functional activity of RLIM/Rnf12 is regulated by phosphorylation-dependent nucleocytoplasmic shuttling. Molecular biology of the cell 24: 3085–3096.

Joazeiro CA, Weissman AM. 2000. RING finger proteins: mediators of ubiquitin ligase activity. Cell 102: 549–552.

Johnsen SA, Gungor C, Prenzel T, Riethdorf S, Riethdorf L, Taniguchi-Ishigaki N, Rau T, Tursun B, Furlow JD, Sauter G et al. 2009. Regulation of estrogen-dependent transcription by the LIM cofactors CLIM and RLIM in breast cancer. Cancer research 69: 128–136.

Kim Y, Bingham N, Sekido R, Parker KL, Lovell-Badge R, Capel B. 2007. Fibroblast growth factor receptor 2 regulates proliferation and Sertoli differentiation during male sex determination. Proceedings of the National Academy of Sciences of the United States of America 104: 16558–16563.

Kniepert A, Groettrup M. 2014. The unique functions of tissue-specific proteasomes. Trends in biochemical sciences 39: 17–24.

Kotaja N, Kimmins S, Brancorsini S, Hentsch D, Vonesch JL, Davidson I, Parvinen M, Sassone-Corsi P. 2004. Preparation, isolation and characterization of stage-specific spermatogenic cells for cellular and molecular analysis. Nature methods 1: 249–254.

Kramer OH, Zhu P, Ostendorff HP, Golebiewski M, Tiefenbach J, Peters MA, Brill B, Groner B, Bach I, Heinzel T et al. 2003. The histone deacetylase inhibitor valproic acid selectively induces proteasomal degradation of HDAC2. The EMBO journal 22: 3411–3420.

Laemmli UK. 1970. Cleavage of structural proteins during the assembly of the head of bacteriophage T4. Nature 227: 680–685.

Margolin G, Khil PP, Kim J, Bellani MA, Camerini-Otero RD. 2014. Integrated transcriptome analysis of mouse spermatogenesis. BMC genomics 15: 39.

Metzger MB, Pruneda JN, Klevit RE, Weissman AM. 2014. RING-type E3 ligases: master manipulators of E2 ubiquitin-conjugating enzymes and ubiquitination. Biochim Biophys Acta 1843: 47–60.

Mikolcevic P, Sigl R, Rauch V, Hess MW, Pfaller K, Barisic M, Pelliniemi LJ, Boesl M, Geley S. 2012. Cyclin-dependent kinase 16/PCTAIRE kinase 1 is activated by cyclin Y and is essential for spermatogenesis. Molecular and cellular biology 32: 868–879.

Mueller JL, Mahadevaiah SK, Park PJ, Warburton PE, Page DC, Turner JM. 2008. The mouse X chromosome is enriched for multicopy testis genes showing postmeiotic expression. Nature genetics 40: 794–799.

Namekawa SH, Park PJ, Zhang LF, Shima JE, McCarrey JR, Griswold MD, Lee JT. 2006. Postmeiotic sex chromatin in the male germline of mice. Current biology: CB 16: 660–667.

O’Donnell L. 2014. Mechanisms of spermiogenesis and spermiation and how they are disturbed. Spermatogenesis 4: e979623.

O’Donnell L, Nicholls PK, O’Bryan MK, McLachlan RI, Stanton PG. 2011. Spermiation: The process of sperm release. Spermatogenesis 1: 14–35.

Oakberg EF. 1956a. A description of spermiogenesis in the mouse and its use in analysis of the cycle of the seminiferous epithelium and germ cell renewal. Am J Anat 99: 391–413.

Oakberg EF. 1956b. Duration of spermatogenesis in the mouse and timing of stages of the cycle of the seminiferous epithelium. Am J Anat 99: 507–516.

Ostendorff HP, Bossenz M, Mincheva A, Copeland NG, Gilbert DJ, Jenkins NA, Lichter P, Bach I. 2000. Functional characterization of the gene encoding RLIM, the corepressor of LIM homeodomain factors. Genomics 69: 120–130.

Ostendorff HP, Peirano RI, Peters MA, Schluter A, Bossenz M, Scheffner M, Bach I. 2002. Ubiquitination-dependent cofactor exchange on LIM homeodomain transcription factors. Nature 416: 99–103.

Ostendorff HP, Tursun B, Cornils K, Schluter A, Drung A, Gungor C, Bach I. 2006. Dynamic expression of LIM cofactors in the developing mouse neural tube. Developmental dynamics: an official publication of the American Association of Anatomists 235: 786–791.

Payer B. 2016. Developmental regulation of X-chromosome inactivation. Seminars in cell & developmental biology 56: 88–99.

Pickart CM. 2001. Mechanisms underlying ubiquitination. Annu Rev Biochem 70: 503–533.

Proudfoot NJ. 2011. Ending the message: poly(A) signals then and now. Genes & development 25: 1770–1782.

Qian X, Mruk DD, Cheng YH, Tang EI, Han D, Lee WM, Wong EW, Cheng CY. 2014. Actin binding proteins, spermatid transport and spermiation. Seminars in cell & developmental biology 30: 75–85.

Rengan AK, Agarwal A, van der Linde M, du Plessis SS. 2012. An investigation of excess residual cytoplasm in human spermatozoa and its distinction from the cytoplasmic droplet. Reprod Biol Endocrinol 10: 92.

Richburg JH, Myers JL, Bratton SB. 2014. The role of E3 ligases in the ubiquitin-dependent regulation of spermatogenesis. Seminars in cell & developmental biology 30: 27–35.

Robinson MD, McCarthy DJ, Smyth GK. 2010. edgeR: a Bioconductor package for differential expression analysis of digital gene expression data. Bioinformatics 26: 139–140.

Robinson MD, Smyth GK. 2007. Moderated statistical tests for assessing differences in tag abundance. Bioinformatics 23: 2881–2887.

Schonhoff SE, Giel-Moloney M, Leiter AB. 2004. Neurogenin 3-expressing progenitor cells in the gastrointestinal tract differentiate into both endocrine and non-endocrine cell types. Developmental biology 270: 443–454.

Sharma U, Conine CC, Shea JM, Boskovic A, Derr AG, Bing XY, Belleannee C, Kucukural A, Serra RW, Sun F et al. 2016. Biogenesis and function of tRNA fragments during sperm maturation and fertilization in mammals. Science 351: 391–396.

Shin J, Bossenz M, Chung Y, Ma H, Byron M, Taniguchi-Ishigaki N, Zhu X, Jiao B, Hall LL, Green MR et al. 2010. Maternal Rnf12/RLIM is required for imprinted X-chromosome inactivation in mice. Nature 467: 977–981.

Shin J, Wallingford MC, Gallant J, Marcho C, Jiao B, Byron M, Bossenz M, Lawrence JB, Jones SN, Mager J et al. 2014. RLIM is dispensable for X-chromosome inactivation in the mouse embryonic epiblast. Nature 511: 86–89.

Sin HS, Ichijima Y, Koh E, Namiki M, Namekawa SH. 2012. Human postmeiotic sex chromatin and its impact on sex chromosome evolution. Genome research 22: 827–836.

Sylwestrak EL, Rajasethupathy P, Wright MA, Jaffe A, Deisseroth K. 2016. Multiplexed Intact-Tissue Transcriptional Analysis at Cellular Resolution. Cell 164: 792–804.

Tian B, Manley JL. 2017. Alternative polyadenylation of mRNA precursors. Nature reviews Molecular cell biology 18: 18–30.

Tomer R, Ye L, Hsueh B, Deisseroth K. 2014. Advanced CLARITY for rapid and high-resolution imaging of intact tissues. Nature protocols 9: 1682–1697.

Trapnell C, Pachter L, Salzberg SL. 2009. TopHat: discovering splice junctions with RNA-Seq. Bioinformatics 25: 1105–1111.

Turner JM. 2007. Meiotic sex chromosome inactivation. Development 134: 1823–1831.

Turner JM. 2015. Meiotic Silencing in Mammals. Annual review of genetics 49: 395–412.

Wang F, McCannell KN, Boskovic A, Zhu X, Shin J, Yu J, Gallant J, Byron M, Lawrence JB, Zhu LJ et al. 2017. Rlim-Dependent and -Independent Pathways for X Chromosome Inactivation in Female ESCs. Cell reports 21: 3691–3699.

Wang F, Shin J, Shea JM, Yu J, Boskovic A, Byron M, Zhu X, Shalek AK, Regev A, Lawrence JB et al. 2016. Regulation of X-linked gene expression during early mouse development by Rlim. Elife 5.

Wang F, Zhao K, Yu S, Xu A, Han W, Mei Y. 2019. RNF12 catalyzes BRF1 ubiquitination and regulates RNA polymerase III-dependent transcription. The Journal of biological chemistry 294: 130–141.

Yamashita M, Honda A, Ogura A, Kashiwabara S, Fukami K, Baba T. 2008. Reduced fertility of mouse epididymal sperm lacking Prss21/Tesp5 is rescued by sperm exposure to uterine microenvironment. Genes Cells 13: 1001–1013.

Yomogida K, Ohtani H, Harigae H, Ito E, Nishimune Y, Engel JD, Yamamoto M. 1994. Developmental stage- and spermatogenic cycle-specific expression of transcription factor GATA-1 in mouse Sertoli cells. Development 120: 1759–1766.

Yoshida S, Takakura A, Ohbo K, Abe K, Wakabayashi J, Yamamoto M, Suda T, Nabeshima Y. 2004. Neurogenin3 delineates the earliest stages of spermatogenesis in the mouse testis. Developmental biology 269: 447–458.

Yuan S, Zheng H, Zheng Z, Yan W. 2013. Proteomic analyses reveal a role of cytoplasmic droplets as an energy source during epididymal sperm maturation. PloS one 8: e77466.

Zheng H, Stratton CJ, Morozumi K, Jin J, Yanagimachi R, Yan W. 2007. Lack of Spem1 causes aberrant cytoplasm removal, sperm deformation, and male infertility. Proceedings of the National Academy of Sciences of the United States of America 104: 6852–6857.

Zhu LJ. 2013. Integrative analysis of ChIP-chip and ChIP-seq dataset. Methods in molecular biology 1067: 105–124.

Zhu LJ, Gazin C, Lawson ND, Pages H, Lin SM, Lapointe DS, Green MR. 2010. ChIPpeakAnno: a Bioconductor package to annotate ChIP-seq and ChIP-chip data. BMC bioinformatics 11: 237.

