## Supplemental Figures for "Deficient spermiogenesis in mice lacking *Rlim*"

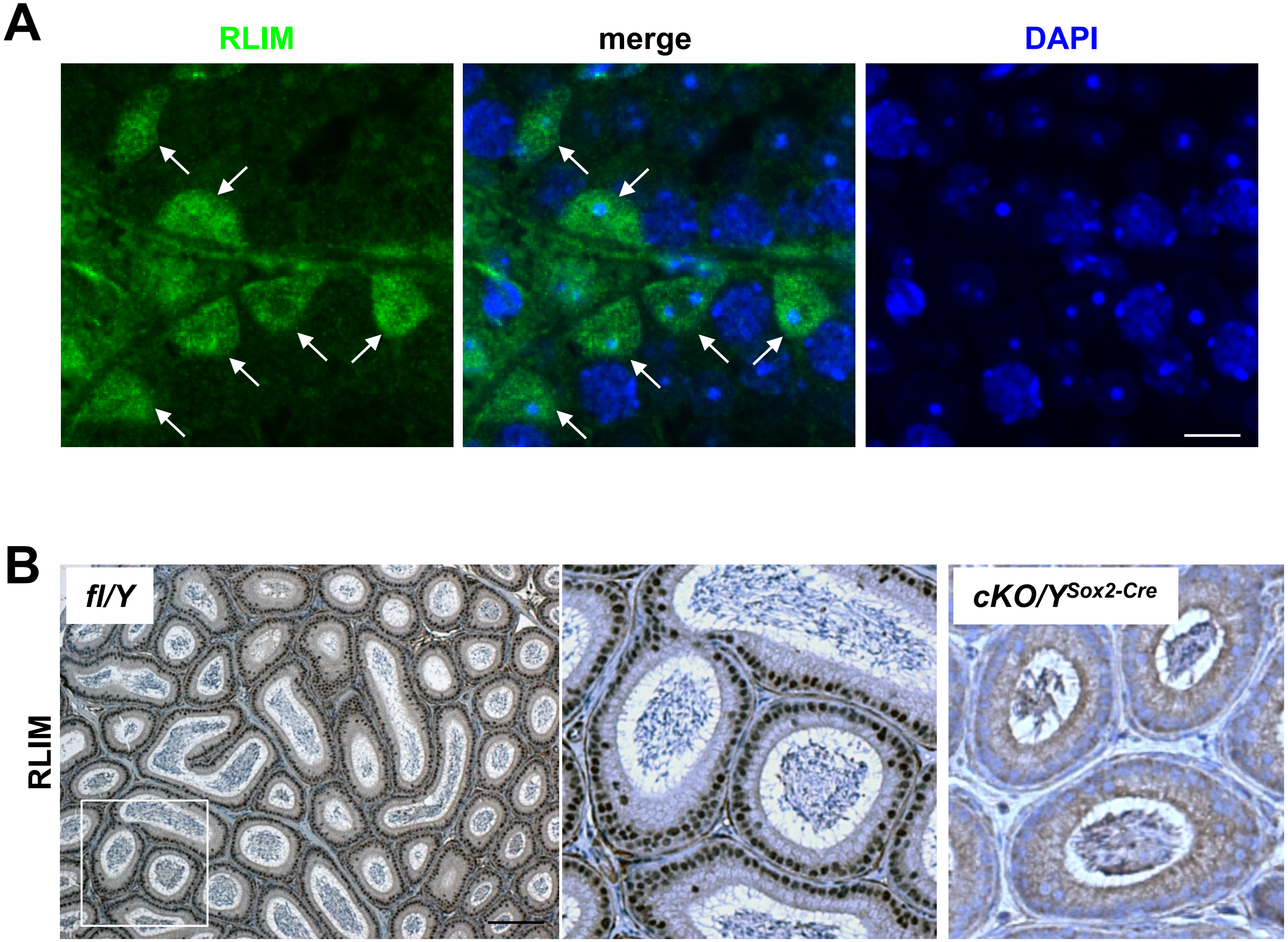
A
RLIM
merge
DAPI
B
fl/Y
cKO/Y^Sox2-Cre
RLIM

**Supplemental figure 1 A)** IHC staining of sections from WT/Y testis using RLIM antibodies. Note RLIM-positive cells in the periphery of seminiferous tubules (arrows) containing triangular-shaped nuclei. Scale bar = 10 µm. **B)** Robust staining of most/all nuclei of epithelial cells lining epididymal tubules in IHC on a cross-section through the Cauda region of fl/Y males using *Rlim* antibodies. cKO/Y sections served as control. Note less sperm in cKO/Y tubules. Scale bar = 50 µm.

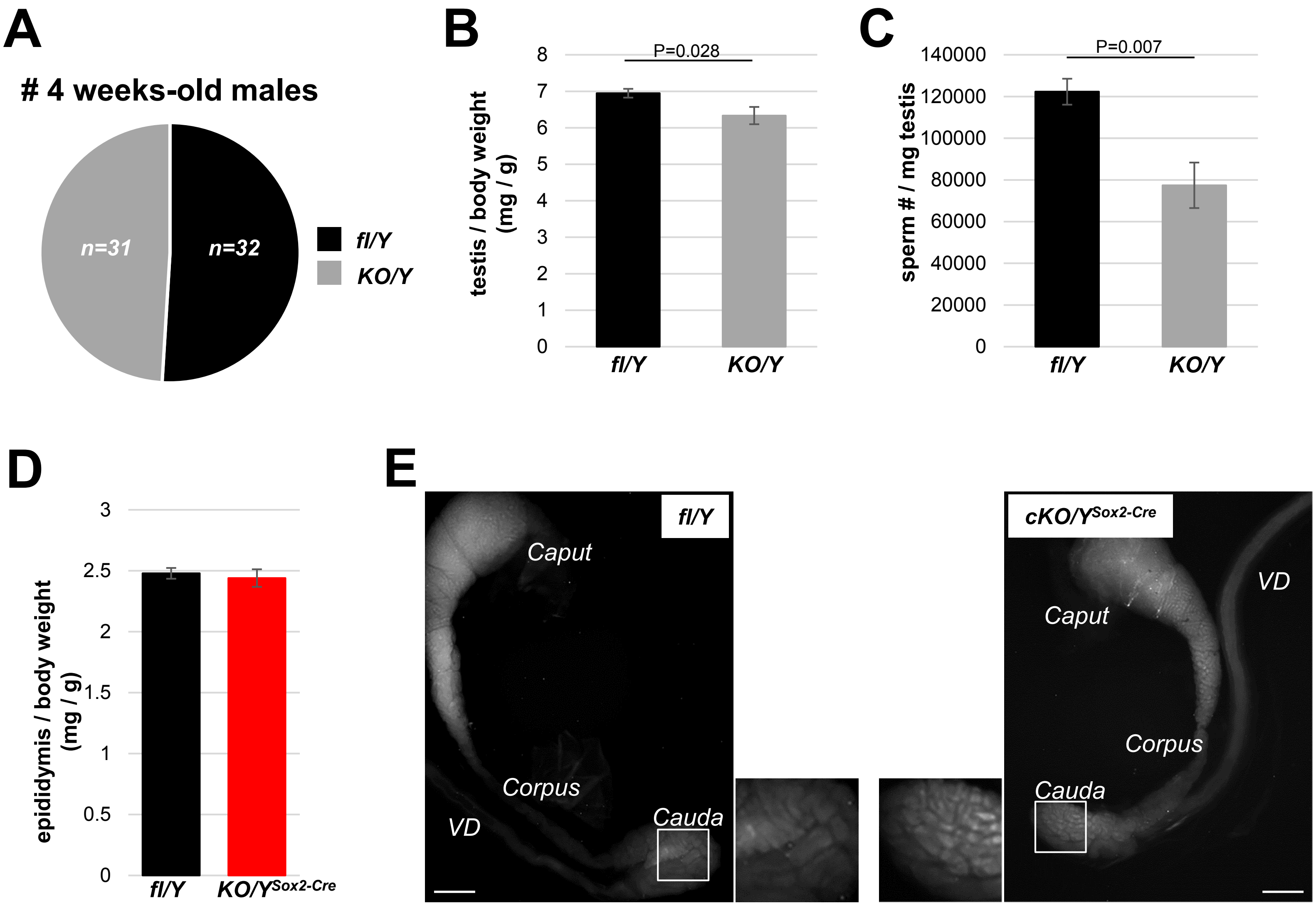
A
### 4 weeks-old males
n=31
n=32
fl/Y
KO/Y
B
P=0.028
testis / body weight (mg / g)
8
7
6
5
4
3
2
1
0
fl/Y
KO/Y
C
P=0.007
sperm # / mg testis
140000
120000
100000
80000
60000
40000
20000
0
fl/Y
KO/Y
D
epididymis / body weight (mg / g)
3
2.5
2
1.5
1
0.5
0
fl/Y
KO/Y^Sox2-Cre
E
fl/Y
Caput
Corpus
Cauda
VD
cKO/Y^Sox2-Cre
VD
Caput
Corpus
Cauda

**Supplemental figure 2** **A)** *Rlim* has no essential functions during male embryogenesis and post-natal development. Using $Rlim^{fl/KO}$ as dames, a similar number of males out of 9 litters receiving a germline *KO* or the *fl Rlim* allele were born and appeared healthy beyond 4 weeks of age. **B)** Significantly decreased weight of testes isolated from 8 weeks old cKO/Y animals (n=8) when compared to fl/Y littermates (n=6). Values were normalized against total body weight and represent the mean ± s.e.m. P values are shown (students t-test). **C)** Significantly decreased numbers of sperm in animals lacking *Rlim*. Cauda epididymal sperm were collected via swim-out in HTF medium. After 10 minutes of swim-out, total sperm numbers were determined (n=6 fl/Y; n=9 cKO/Y). s.e.m. and P values are indicated. **D)** Similar weights of fl/Y and cKO/$Y^{Sox2\text{-}Cre}$ epididymides (n=9 animals, each). **E)** Visualization of epididymal structure via de-lipidation. Caput, Corpus, Cauda regions as well as Vas Deferens (VD) are indicated. Note that mice lacking *Rlim* display a slightly thinner tubular structure within the Cauda, consistent with a lower sperm content.

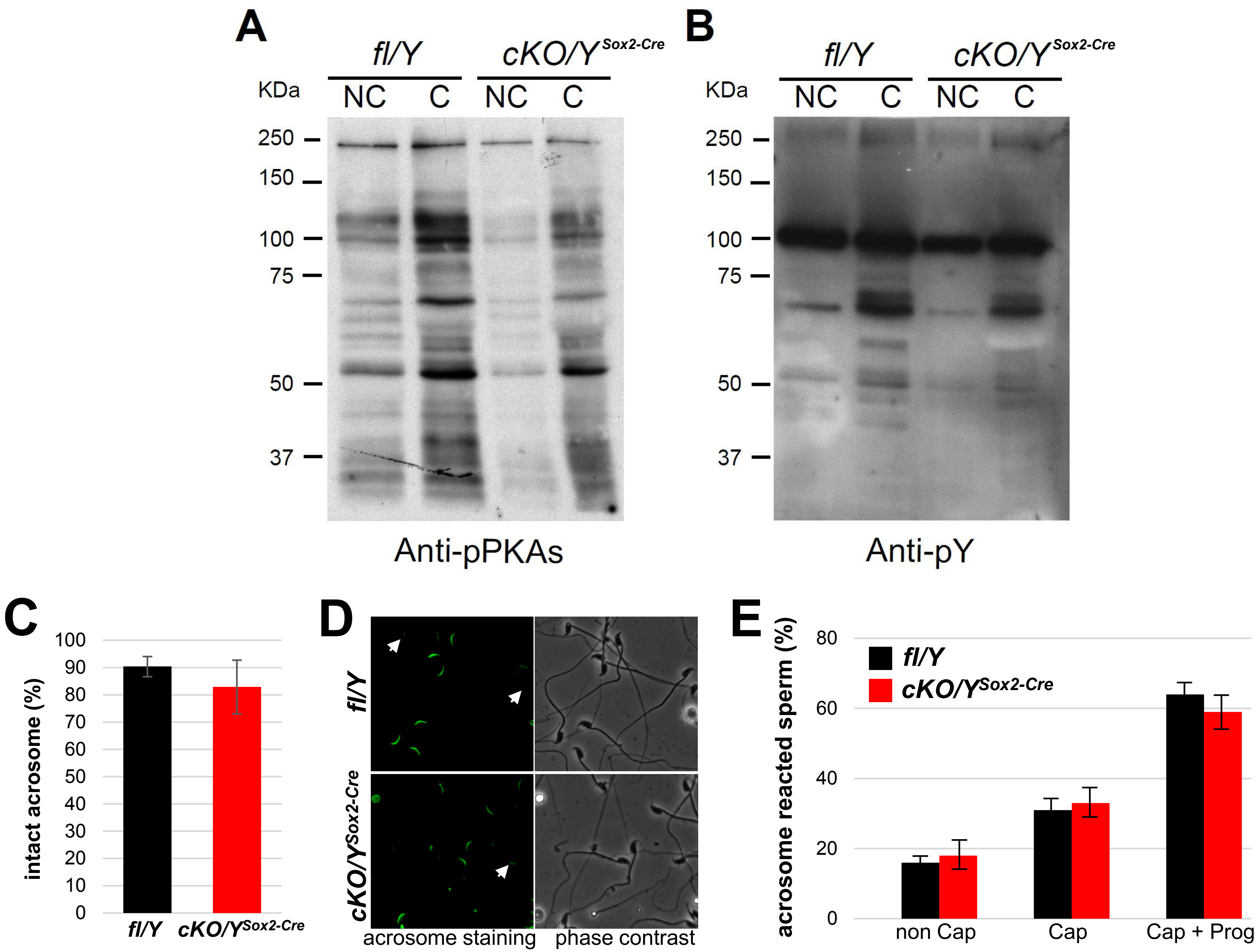
A
fl/Y
cKO/Y Sox2-Cre
KDa
NC C NC C
250
150
100
75
50
37
Anti-pPKAs
B
fl/Y
cKO/Y Sox2-Cre
KDa
NC C NC C
250
150
100
75
50
37
Anti-pY
C
intact acrosome (%)
100
90
80
70
60
50
40
30
20
10
0
fl/Y
cKO/Y Sox2-Cre
D
fl/Y
cKO/Y Sox2-Cre
acrosome staining
phase contrast
E
acrosome reacted sperm (%)
80
60
40
20
0
fl/Y
cKO/Y Sox2-Cre
non Cap
Cap
Cap + Prog

**Supplemental figure 3** Evaluation of molecular and physiological events related to sperm capacitation. Sperm were incubated at 37 °C for 60 min in m-TYH (non-capacitating conditions, NC) or in m-TYH supplemented with 15 mM NaHCO3 and 5 mg/ml BSA (capacitating conditions, C). **A)** After 60 min of incubation proteins were extracted, subjected to SDS–PAGE, and electro-transferred to PVDF membranes. Then, phosphorylation of PKA substrates was detected by immunoblotting with Anti-pPKAs antibody. Representative image of four independent experiments. **B)** Membranes were stripped and then phosphorylation of proteins in tyrosine residues (pY) was detected by immunoblotting with Anti-pY antibody. Representative image of four independent experiments. **C)** Evaluation of capacitation through the ability to acrosome react. After 60 min of capacitation in m-TYH Cap medium, sperm samples were incubated with progesterone (Prog, 10 μM) or with DMSO (vehicle) in m-TYH Cap at 37 °C for additional 30 min, and then acrosomal status was assessed by Alexa Fluor 488-PNA staining. Acrosome reaction after 90 min in non-capacitating (Non Cap) conditions was also evaluated. Results are indicated in percentage of acrosome reacted sperm. n=4, at least 200 sperm per sample were counted. Values represent the mean ± s.e.m. **D)** Evaluation of acrosomal status by Alexa Fluor 488-PNA staining. Representative fluorescence images of the acrosomal staining (left panel), and their corresponding phase contrast images (right panel). **E)** Sperm displaying uniform fluorescence along their acrosomes were counted as intact. n=9, at least 200 sperm per sample were counted. Values represent the mean ± s.e.m.

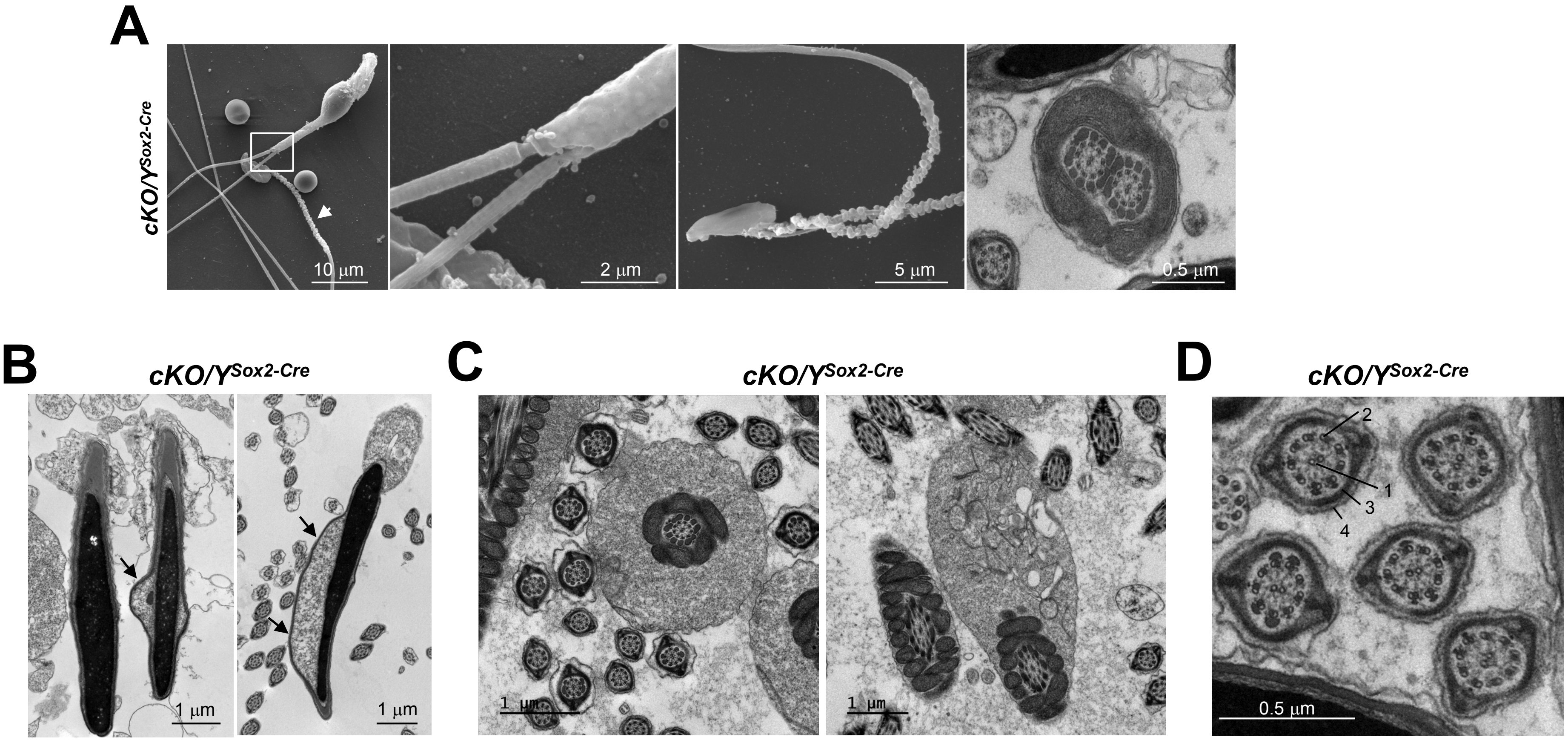
A
cKO/Y^Sox2-Cre
10 μm
2 μm
5 μm
0.5 μm
B
cKO/Y^Sox2-Cre
1 μm
1 μm
C
cKO/Y^Sox2-Cre
1 μm
1 μm
D
cKO/Y^Sox2-Cre
2
1
3
4
0.5 μm

**Supplemental figure 4** Evaluation of sperm structure within testes of cKO/$Y^{Sox2\text{-}Cre}$ animals. **A)** Examples of cKO/$Y^{Sox2\text{-}Cre}$ sperm exhibiting two axonemes (SEM). Left panel: Sperm with two axonemes. Boxed area is shown in higher magnification. Middle panel: Sperm with two axonemes and ruptured midpiece. Right panel: TEM showing sperm with two axonemes, as indicated by the same orientation of dynein arms. **B)** Head malformations as detected in TEM on testes sections are indicated by arrows. Arrows point at cytoplasmic pockets. **C, D)** No major structural deficiencies were detected via TEM on testes sections in the midpiece (C) and the tail (D). 1=inner microtubule doublet of the axoneme (2); 2=outer microtubule doublet of the axoneme (9); 3=fibrous sheath; 4=plasma membrane.

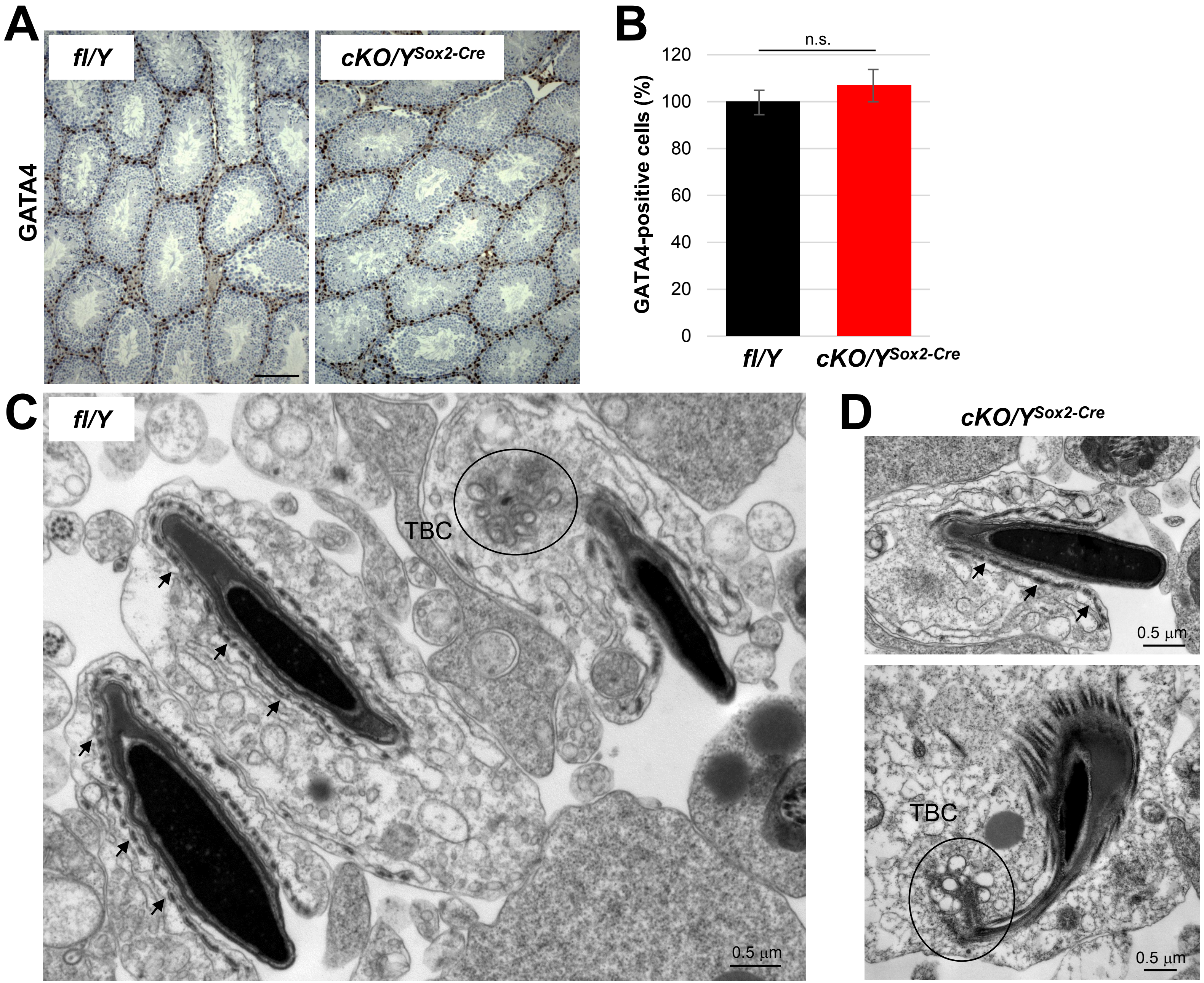

A
fl/Y
cKO/YSox2-Cre
GATA4
B
n.s.
GATA4-positive cells (%)
120
100
80
60
40
20
0
fl/Y
cKO/YSox2-Cre
C
fl/Y
TBC
0.5 μm
D
cKO/YSox2-Cre
0.5 μm
TBC
0.5 μm

**Supplemental figure 5** Normal numbers of Sertoli cells in testes lacking *Rlim*. **A)** DAB staining of paraffin-embedded testes sections of cKO/$Y^{Sox2\text{-}Cre}$ and fl/Y animals using antibodies against Sertoli cell marker GATA4. **B)** Quantification of GATA4-positive cells within seminiferous tubules. **C, D)** Normal development of Sertoli cell structures during spermiation in animals lacking *Rlim* as shown via TEM on testes sections including the apical ectoplasmic specialization (ES; arrows) and the apical tubulobulbar complex (TBC circled).
